# Similar fertility losses in response to heat are followed by distinct recovery trajectories across seven *Drosophila* species

**DOI:** 10.64898/2026.10.04.756584

**Authors:** Wayne Wen-Yeu Wang, Shelby L. Jackson, Alexis F. Bartek, Colin D. Meiklejohn, Kristi L. Montooth

**Affiliations:** School of Biological Sciences, University of Nebraska-Lincoln

**Keywords:** Climate change, Thermal physiology, Thermal fertility limits, Sperm storage

## Abstract

Heat-induced sterility is widespread across ectotherm species, yet its underlying mechanisms remain poorly understood. In mated females, thermal stress can disrupt female reproductive physiology while also affecting sperm stored within the reproductive tract, making the mechanisms of fertility loss difficult to distinguish. Here, we investigated the extent and recovery dynamics of female heat-induced fertility loss across seven *Drosophila* species. We tracked fertility after acute heat exposure, tested whether remating restored fertility, quantified stored sperm in two closely related species, and examined whether reproductive thermal sensitivity covaried with adult heat tolerance. We found that heat exposure reduced egg-hatching success in all seven species, but recovery differed among species. Fertility recovered within several days in most species, whereas *D. simulans* showed persistent declines and *D. melanogaster* showed limited recovery. Remating restored or maintained fertility in both species. Stored sperm declined through time in both *D. simulans* and *D. mauritiana*, but *D. mauritiana* retained more sperm at later sampling points, consistent with their contrasting patterns of fertility decline and maintenance. Across species, greater adult heat tolerance was associated with smaller immediate reductions in fertility, but not with cumulative reproductive loss across subsequent days. Together, these findings show that similar initial fertility losses can lead to markedly different recovery trajectories and highlight the importance of comparative, multi-species approaches that integrate female physiology, sperm-storage dynamics, mating behavior, and whole-organism thermal tolerance when assessing reproductive resilience to climate warming.

## Introduction

Reproduction is fundamental to population persistence and is often among the first biological processes disrupted by environmental change (Parratt et al., 2021; van Heerwaarden and Sgrò, 2021). Under climate warming, extreme heat can impair reproduction at temperatures well below lethal limits, constraining population persistence even when adults survive acute heat events (Review in Wang and Gunderson, 2022; Walsh et al., 2019; Snook et al., 2026; Dougherty et al., 2024). Understanding reproductive thermal sensitivity is therefore essential for predicting how organisms will respond to increasingly frequent and intense thermal extremes (Parratt et al., 2021; van Heerwaarden and Sgrò, 2021).

Fertility loss has also frequently been observed when females experience elevated temperatures (McAfee et al., 2020; Walsh et al., 2022; Parratt et al., 2021). While male reproductive function is often particularly sensitive to heat stress (Dougherty et al., 2024; Iossa, 2019; David et al., 2005), female reproductive responses to heat stress have been comparatively understudied. This is an important gap because successful reproduction depends on multiple processes within females that may be thermally sensitive in ectotherms, including oogenesis, oocyte maturation, sperm storage, fertilization, and the completion of embryonic development (Walsh et al., 2019, 2022; Snook et al., 2026; McAfee et al., 2020). Heat exposure could therefore reduce female reproductive success through several distinct pathways, each with different consequences for the duration and reversibility of fertility loss. This calls for an empirical approach that integrates across multiple female and male processes to fully understand the thermal sensitivity of reproduction.

Stored sperm is one of several reproductive components vulnerable to heat stress, but its role in heat-induced sterility has only recently been recognized (McAfee et al., 2020; Walsh et al., 2022; Sales et al., 2018; Wang and Gunderson, 2022). In species with prolonged sperm storage, thermal stress can damage sperm within female sperm-storage organs, causing fertility loss that persists after the heat event (McAfee et al., 2020; Walsh et al., 2022; Sales et al., 2018). For example, reduced stored-sperm viability following thermal extremes has been associated with reproductive failure in queen honeybees (McAfee et al., 2020). If persistent fertility loss is associated primarily with damaged or limiting stored sperm, females should show little recovery without additional sperm but regain fertility after remating (Walsh et al., 2022; Meena et al., 2025; Sales et al., 2018).

In contrast, if heat causes transient disruption of female reproductive physiology or developing oocytes, fertility may recover through time without remating (David et al., 2005; Meiselman et al., 2018). Therefore, following fertility through time and examining responses to remating can help distinguish between these mechanisms of heat-induced fertility loss.

*Drosophilid* flies provide a powerful system for testing these predictions and have long served as models of thermal physiology. Their short generation times, well-established culturing methods, and genetic tools, including fluorescent-labelled sperm, allow direct examination of how different components of reproduction respond to heat stress. Although heat-induced male sterility is widespread across *Drosophilids*, studies of female heat sensitivity have often focused on unmated females to isolate direct effects on female reproduction (David et al., 2005; Meena et al., 2024; van Heerwaarden and Sgrò, 2021). Fewer studies have examined mated females, but the results suggest that stored sperm and other post-mating reproductive processes may be particularly vulnerable to heat stress, often leading to prolonged reductions in fertility (*D. melanogaster*: Meena et al., 2025; Bundgaard and Barker, 2017; *D. Virilis* and *Zaprionus indianus*: Walsh et al., 2022). Whether similar heat-induced fertility losses arise through comparable reproductive processes across species, and whether species differ in their capacity to recover remains unknown.

Reproductive thermal sensitivity may co-evolve with whole-organism thermal physiology (Parratt et al., 2021; van Heerwaarden and Sgrò, 2021; Wang and Gunderson, 2022). Because reproduction contributes directly to fitness, species adapted to warmer environments may evolve both greater adult heat tolerance and more heat-resistant reproductive processes. Under this thermal co-adaptation hypothesis, species with higher adult heat tolerance should experience smaller reproductive losses following heat exposure (Angilletta, 2009; Angilletta et al., 2006). Such a pattern would suggest that reproductive physiology can adapt to warmer environments rather than acting as a fixed thermal constraint. Conversely, weak correspondence between these traits would suggest that reproductive and whole-organism thermal physiology are shaped partly independently.

Here, we investigated the extent and recovery dynamics of heat-induced fertility loss in mated females across seven *Drosophila* species. We quantified egg production, hatch rate, and cumulative reproductive output following acute heat exposure to determine how both immediate and longer-term reproductive costs varied among species (Fig. 1A). We then tested whether a subsequent mating opportunity restored fertility in species showing persistent reproductive impairment following female heat exposure (Fig. 1 A). We predicted that fertility associated with heat-induced sperm loss would improve after females received a new ejaculate, whereas transient disruption of female reproductive processes would recover without remating and indicate thermal tolerance of stored sperm. To examine whether sperm-storage dynamics contributed to differences in fertility maintenance, we quantified fluorescently labelled sperm within the seminal receptacles of *D. simulans* and *D. mauritiana*, two closely related species with contrasting fertility trajectories through time (Fig. 1B). Finally, using a phylogenetic comparative approach, we tested whether adult heat tolerance (CTmax) was associated with either the immediate thermal sensitivity of fertility or cumulative reproductive loss. Together, these experiments provide an integrative framework for understanding how reproductive systems respond to thermal stress.

**Fig. 1.**
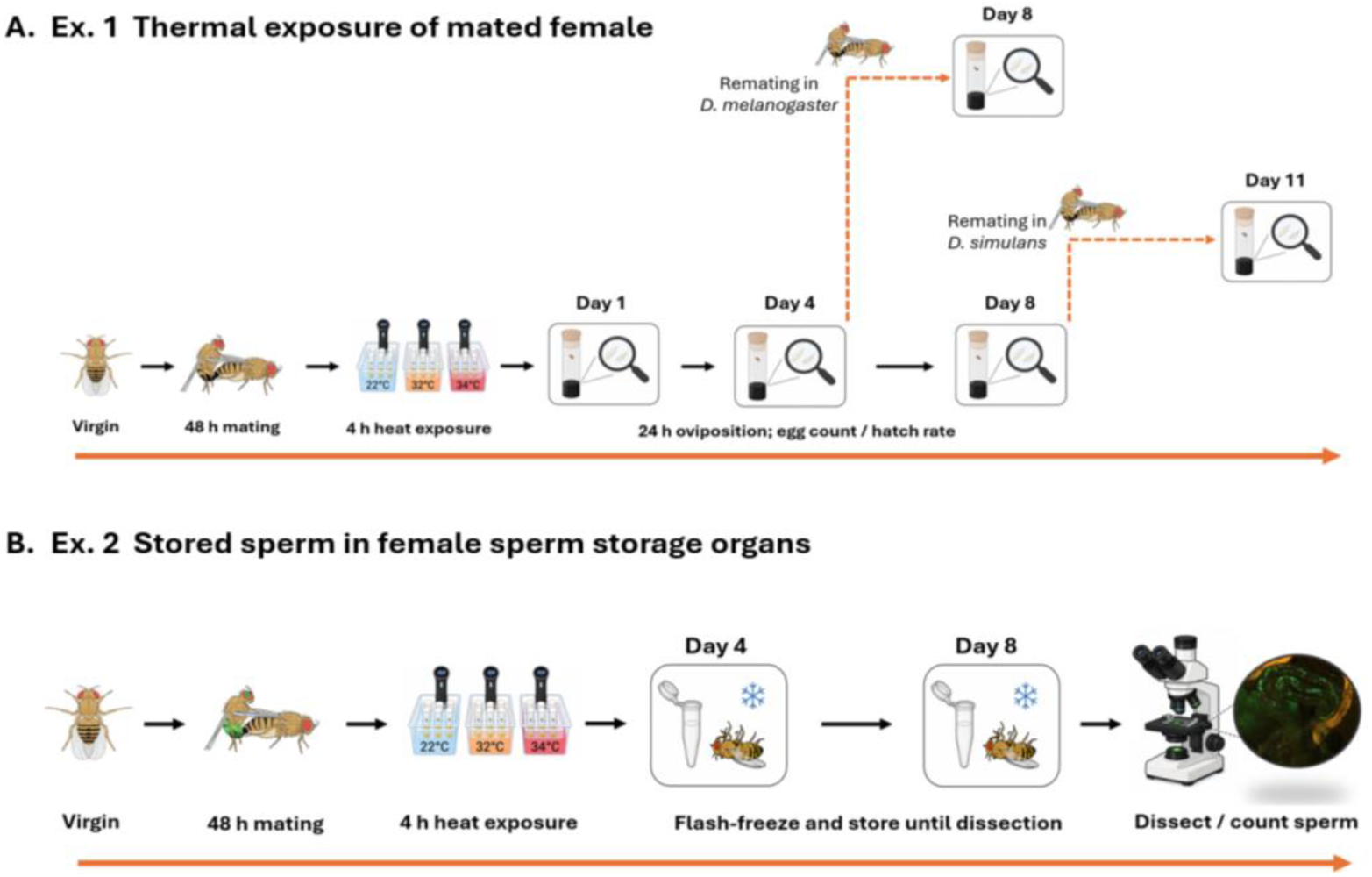
Overview of the experimental design. **(A)** Mated females were exposed to 22, 32, or 34°C for 4 h, and egg number and hatching rate were measured through time. Subset of females from the two species showing prolonged fertility loss were later remated with untreated males and re-assayed. **(B)** Wild-type *D. simulans* (NS40) females were mated with Protamine B–GFP *D. simulans males*, and wild-type *D. mauritiana* females were mated with RFP-labelled *D. mauritiana* males. Females were then exposed to heat, flash-frozen at designated time points, and dissected to quantify fluorescently labelled sperm in the seminal receptacle.

## Methods

### Study organisms

We tested seven species of *Drosophila*: *D. melanogaster*, *D. simulans*, *D. mauritiana*, *D. sechellia*, *D. yakuba*, *D. santomea*, and *D. teissieri* (Fig. 2). The *D. melanogaster* population was collected near Lincoln, Nebraska, USA, in 2016 and has been maintained as an outbred laboratory population. The remaining species were maintained as laboratory stocks: *D. simulans* NS40, *D. mauritiana* (DSSC 14021-0241.05), *D. sechellia* (DSSC 14021-0248.25), *D. yakuba* (DSSC 14021-0261.40), *D. santomea* (14021-0248.01), and *D. teissieri* (DSSC 14021-0257.01-J90001).

**Fig. 2.**
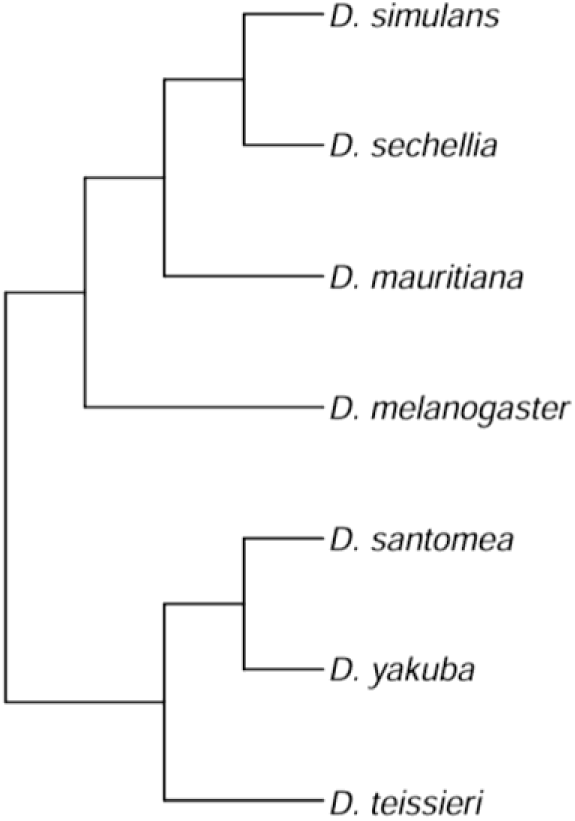
Phylogenetic relationship of the seven species studied.

In Experiment 2, males from a Protamine B–GFP *D. simulans* line (w+; pBac{3xP3-EGFP, ProtB-EGFP}) and an RFP-labelled *D. mauritiana* line (w; P{w8, ProtB-DsRed-monomer, w+}; obtained from John Belote) were used to quantify sperm within the female reproductive tract. All fly stocks were maintained on cornmeal-molasses-Torula yeast diet at 22°C under a 12:12 light:dark photoperiod.

### Experiment 1: Fertility following thermal exposure of mated females

To investigate the effects of heat exposure on female fertility, mated females were exposed 48 hours after mating to a heat treatment for 4 hours, and their fertility was monitored through time (Fig. 1 A).Virgin flies of both sexes were collected upon eclosion and housed in single-sex groups of fewer than 30 flies per food vial. Three to four days after collection, females were paired with a single male using an aspirator. Each pair was housed together for 48 h, after which females were separated from males and transferred to fresh vials in preparation for thermal exposure. The original mating vials were checked for the presence of larvae to confirm that mating occurred. Females whose mating vials showed no evidence of larval development were excluded from the experiment. Mated females were randomly assigned to one of three temperature treatments (22°C, 32°C, or 34°C) for four hours by placing vials in water baths such that the entire vial was submerged below the water line (Fig. 1 A). The 34°C treatment was only used for *D. melanogaster* and *D. simulans,* because this temperature caused high mortality in other species. Water temperature was maintained using precision immersion circulators (BLITZHOME), with separate water baths for each temperature treatment. Following thermal exposure, females were transferred to fresh vials using an aspirator and vials that housed the females during heat exposure were discarded. This assured that all eggs scored for hatching were laid at 22°C. Fertility was measured on the day of exposure (Day 1), and again 4 and 8 days after heat exposure. Multiple species were tested concurrently throughout the experimental period, with individuals from different species interspersed across treatments and experimental dates to minimize potential temporal or batch effects. Sample sizes varied among species, temperature treatments, and sampling days; detailed sample sizes are provided in Table S1.

### Fertility measurements

To quantify fertility, females were transferred to food vials with black food dye to facilitate egg counting. Females were allowed to oviposit for 24 h before being transferred to a new vial. Experimental vials were then maintained at room temperature for an additional 24 to 48 hours before being scored under a dissecting microscope. For each vial, the total number of eggs and the number of hatched eggs were recorded. Hatched eggs were identified by the presence of a collapsed eggshell or/and an open chorion. Hatch rate was calculated as the proportion of scored eggs that hatched. In addition to day-specific measurements, cumulative reproductive output was quantified as the total number of hatched eggs produced by each female across Days 1, 4, and 8.

### Remating experiment

To determine whether an additional mating opportunity could restore fertility following heat exposure, a subset of *D. simulans* and *D. melanogaster* females were paired with a new 3-day-old virgin male for 48 hours before the male was removed. In *D. melanogaster*, females were provided with a remating opportunity after the Day 4 fertility assessment and fertility was measured again on Day 8. In *D. simulans*, the remating opportunity was provided after the Day 8 assessment and fertility was measured again on Day 11.

### Experiment 2: Stored sperm in female sperm storage organs after heat exposure

To test whether differences in fertility recovery between *D. simulans* and *D. mauritiana* were associated with changes in stored sperm, we quantified sperm remaining in the female seminal receptacle (SR) following thermal exposure (Fig. 1 B). Virgin wild-type females of each species were collected upon eclosion and paired with males carrying fluorescently labelled sperm; NS40 *D. simulans* females were paired with Protamine B– GFP *D. simulans* males, whereas wild-type *D. mauritiana* females were paired with Protamine B-RFP-labelled *D. mauritiana* males. Flies were paired at 3–4 days of age and maintained with males for 48 h following the mating protocol described in Experiment 1. After mating, *D. simulans* females were randomly assigned to 22, 32, or 34°C treatments, whereas *D. mauritiana* females were exposed only to 22 or 32°C because 34°C caused high mortality in this species. All thermal exposures lasted 4 hours and followed the procedures described above in experiment 1. Females were flash-frozen on Days 1, 4 and 8 following heat exposure and stored until dissection. We counted fluorescently labelled sperm manually under a fluorescence microscope. Although females also store sperm in the paired spermathecae, these organs were not included because their compact, three-dimensional structure prevented reliable counts of individual sperm. Detailed sample sizes for each species, temperature treatment, and sampling day are in Table S8.

### Data analysis

All analyses were conducted in R version 4.4.3 (R Core Team, 2025). For each species, hatch rate and total number of eggs laid per female were analyzed using linear mixed-effects models in the lme4 and lmerTest packages (Bates et al., 2015; Kuznetsova et al., 2017). Models included temperature, sampling day, and their interaction as fixed effects, with female identity as a random intercept to account for repeated measurements (response ∼ temperature * day + (1 | female)). For hatch rate, observations with fewer than five scored eggs were excluded, whereas all egg-count observations were retained for analyses of egg production. Type III ANOVA tests evaluated the effects of temperature, day, and their interaction. Estimated marginal means and pairwise contrasts were obtained using emmeans (Lenth, 2024), with Tukey-adjusted *P*-values for comparisons among temperatures within sampling days and among days within temperatures. For cumulative reproductive output, we combined the number of hatched eggs (offspring) for each female across Days 1, 4, and 8. Cumulative output was analyzed separately for each species, and pairwise differences among temperatures were evaluated using Welch’s *t*-tests. For species with three temperature treatments, *P*-values from the three pairwise comparisons were adjusted within species using the Holm method to account for multiple testing. For the analysis of cumulative reproductive output, we excluded *D. melanogaster* females assigned to the remating treatment because the remating opportunity occurred before the Day 8 measurement, whereas *D. simulans* females assigned to the remating treatment were retained because the remating opportunity occurred only after Day 8.

Remating effects were analyzed separately for *D. melanogaster* and *D. simulans* for both hatch rate and egg production. In *D. melanogaster*, remating occurred after Day 4 and responses were compared between Days 4 and 8; in *D. simulans*, remating occurred after Day 8 and responses were compared between Days 8 and 11 (Fig. 1A). Models included temperature, sampling day, remating status, and their interactions as fixed effects, with female identity as a random intercept. Pairwise contrasts compared remated and non-remated females and changes through time within each temperature-by-remating group.

To quantify changes in stored sperm abundance, we counted the number of sperm remaining in the seminal receptacle (SR) of *D. simulans* and *D. mauritiana* females at each sampling day. Because sperm counts were strongly right-skewed, sperm number was log-transformed as log10 (sperm number + 1) prior to analysis. For each species, the effects of temperature treatment, sampling day, and their interaction on SR sperm number were evaluated using linear models. Estimated marginal means and pairwise comparisons were obtained using the “emmeans” package (Lenth, 2024), with Tukey-adjusted *P*-values for post hoc tests. To compare sperm retention between species, data from the shared 22 and 32°C treatments were analyzed using a model including species, temperature, and sampling day.

To test whether female fertility thermal sensitivity was associated with adult heat tolerance across species, we quantified thermal sensitivity in two ways. Immediate fertility sensitivity was calculated as the difference in Day 1 hatch rate between 32 and 22°C (32°C − 22°C), such that more negative values indicated greater heat-induced fertility loss. Day 1 was used because it captured the immediate response to heat exposure before substantial recovery occurred. We also quantified cumulative reproductive sensitivity as the proportional change in cumulative offspring production across Days 1, 4, and 8 between 32 and 22°C, calculated as (32°C − 22°C)/22°C, with more negative values indicating greater cumulative reproductive loss. Species-specific CTmax values were obtained from Parratt et al. (2021). Associations between each fertility metric and CTmax were tested separately using phylogenetic generalized least squares (PGLS), with Pagel’s λ estimated by maximum likelihood.

## Results

### Cumulative reproductive costs of heat exposure

Heat exposure reduced the cumulative number of hatched offspring in most species (Fig. 3; Supplementary Table S1). Between 22 and 32°C, *D. simulans*, *D. santomea*, *D. yakuba*, and *D. teissieri* showed significant declines, ranging from approximately 40% in *D. yakuba* to 77% in *D. teissieri*. In *D. simulans*, output declined further at 34°C, reaching approximately 71% below 22°C (Fig. 3; Table S1; *P* = 0.006). In *D. melanogaster*, the reduction at 32°C was not significant, but cumulative offspring production declined by approximately 53% at 34°C relative to 22°C. No significant reduction was detected in *D. sechellia* or *D. mauritiana.* These results suggest that short-term heat exposure can translate into substantial losses in reproductive output. The magnitude of these losses also varied among species.

**Fig. 3.**
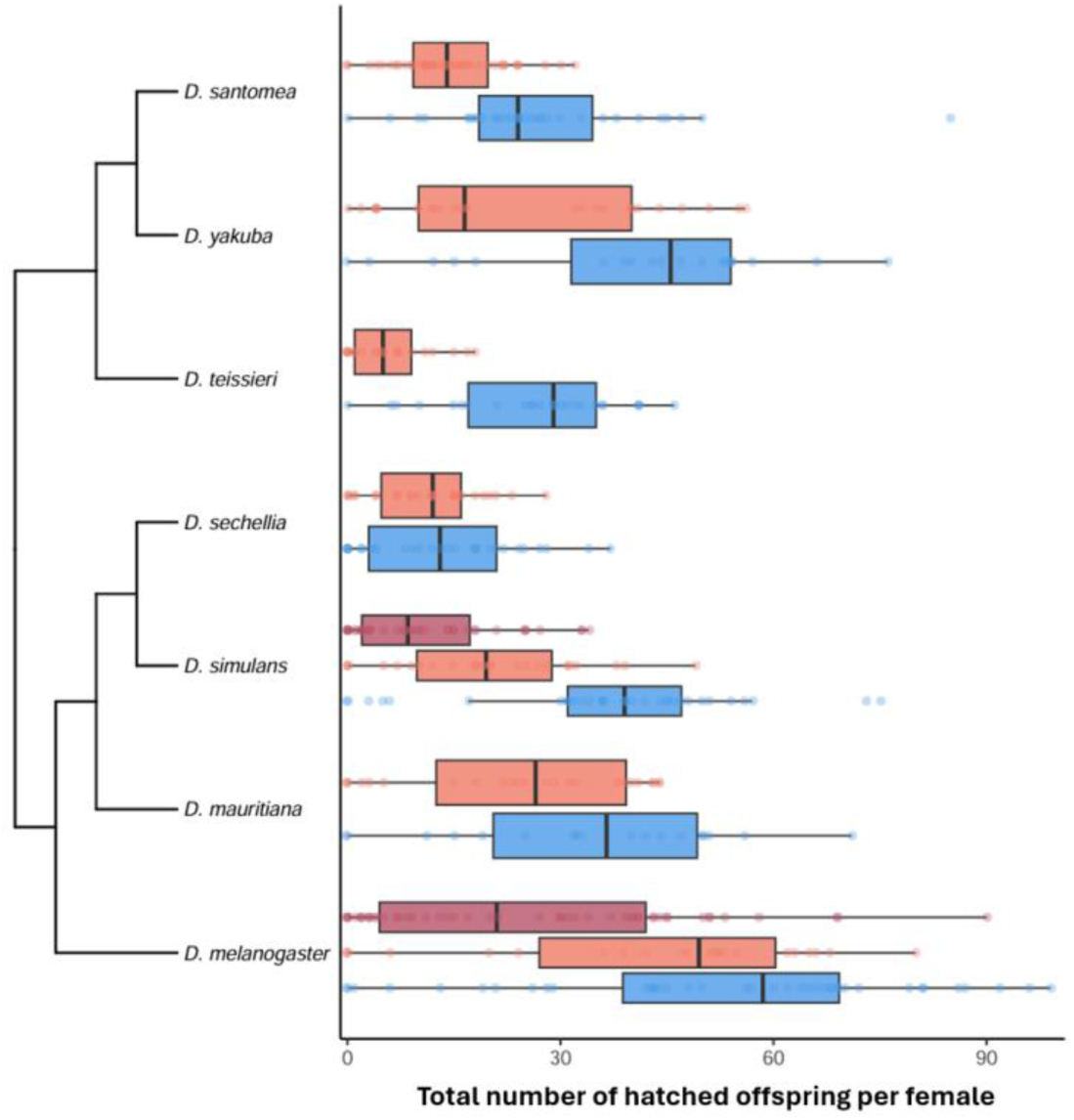
Cumulative reproductive output following heat exposure across seven *Drosophila* species. Total number of hatched offspring produced by each female across all days following exposure to 22, 32, or 34°C. The 34°C treatment was included only for *D. melanogaster* and *D. simulans*. Boxes show the median and interquartile range, whiskers extend to 1.5 × the interquartile range, and points represent individual females. The phylogeny indicates relationships among species.

### Fertility loss and recovery following heat exposure

Heat exposure immediately reduced the hatch rate across all species, although the magnitude varied among species (Fig. 4; Tables S2, S3). On Day 1, hatch rate was significantly lower at 32°C in *D. simulans*, *D. sechellia*, *D. mauritiana*, *D. santomea*, *D. yakuba*, and *D. teissieri*, with absolute reductions in hatch rate ranging from 0.30 to 0.66 relative to 22°C (Fig. 4; Tables S3; all *P* ≤ 0.003). In *D. melanogaster*, the reduction was significant only at 34°C (Fig. 4 B; Tables S3; *P* = 0.012). In *D. simulans*, the effect increased with temperature, with hatch rate declining significantly between 22°C and 32°C and between 32C and 34°C on Day 1 (Fig. 4 A; Tables S3; all pairwise *P* ≤ 0.002).

**Fig. 4.**
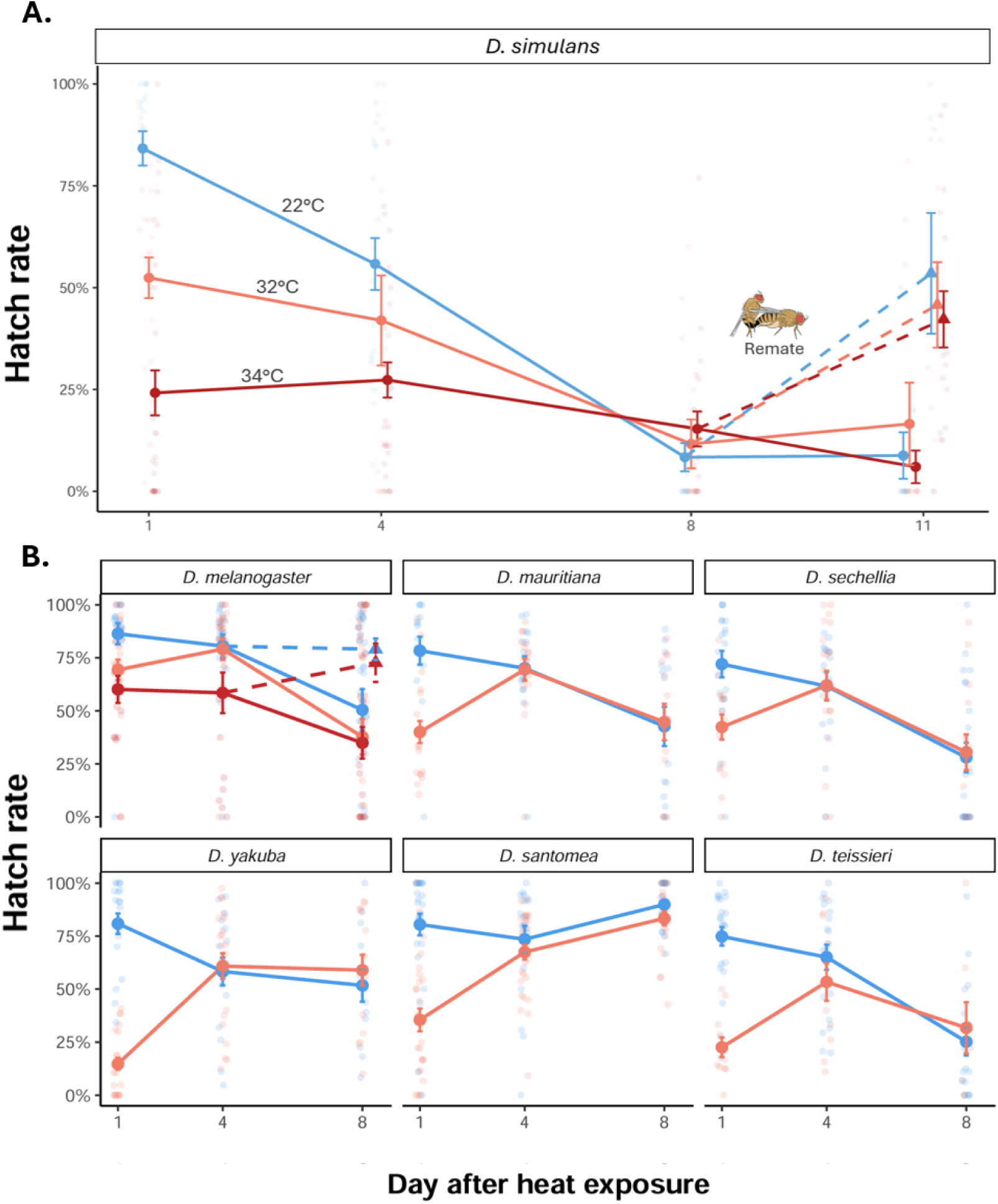
Recovery of female fertility following heat exposure in Drosophila. **(A)** Hatch rate of mated *D. simulans* females exposed to 22°C (blue), 34°C (red) or 32°C (orange). Hatch rate was assessed on Days 1, 4, and 8 following heat exposure. A subset of females was remated after Day 8 and reassessed on Day 11 (dashed lines). **(B)** Hatch rate of mated females from six other *Drosophila* species following exposure to 22°C (blue), 32°C (orange), or 34°C (red; *D. melanogaster* only). Points represent model-estimated means ± 1 SE, and transparent points show individual females. In *D. melanogaster*, triangle and dashed lines indicate females remated after Day 4, whereas filled circles and solid lines represent non-remated females.

By Day 4 and without remating, temperature differences had disappeared in five of these species, with hatch rate at 32°C increasing by approximately 0.31–0.46 from Day 1 to Day 4 in *D. mauritiana*, *D. santomea*, *D. yakuba*, and *D. teissieri* (Fig. 4; Tables S4; all *P* ≤ 0.014). These recoveries suggest that the effects of heat on fertility were largely transient in these species and were not caused by loss of stored sperm due to heat. In contrast, hatch rate in *D. simulans* declined from Day 1 to Day 4 at 32°C (Fig. 4 A; Tables S4; 0.491 ± 0.050 to 0.305 ± 0.075, *P* = 0.028) and declined further by Day 8 (0.084 ± 0.042, *P* = 0.017), while no significant change occurred between Days 1 and 4 at 34°C (Tables S4; *P* = 0.399). Similarly, hatch rate in *D. melanogaster* showed no significant recovery between Days 1 and 4 at 34°C (Fig. 4 B; Tables S4; 0.601 ± 0.063 to 0.585 ± 0.096, *P* = 0.995). These more persistent effects of heat exposure suggest longer-lasting damage to developing oocytes or stored sperm.

### Fertility declines through time independent of temperature

Hatch rate also declined through time regardless of temperature, but the magnitude of decline varied among species (Fig. 4; Tables S4). The strongest decline occurred in *D. simulans*: at 22°C, observed mean hatch rate fell from 0.814 ± 0.044 on Day 1 to 0.100 ± 0.036 on Day 8 (Tables S4; *P* < 0.001), and at 32°C from 0.491 ± 0.050 to 0.084 ± 0.042 (*P* < 0.001). In comparison, hatch rate in *D. mauritiana* declined more moderately at 22°C, from 0.797 ± 0.059 to 0.427 ± 0.093 (Tables S4; *P* < 0.001), and showed no significant decline at 32°C (*P* = 0.297). Among the remaining species that declined over time at 22°C, reductions in hatch rate from Day 1 to Day 8 ranged from approximately 0.29 to 0.50. In contrast, hatch rate in *D. santomea* remained high through time at 22°C (Days 1: 0.805 ± 0.051, Days 4: 0.735 ± 0.063, and Days 8: 0.899 ± 0.020; Tables S4; all *P* ≥ 0.073). This pattern suggests a natural decline in fertility through time that is largely independent of temperature and may reflect species differences in the ability to maintain sufficient stored sperm.

### Re-mating rescues fertility in *D. simulans* and *D. melanogaster*

We next tested whether a new mating opportunity could restore fertility in *D. simulans* and *D. melanogaster*. In *D. simulans*, females were allowed to remate after Day 8, and hatch rate increased significantly by Day 11 in females that remated (Fig 4A; Tables S5), from 0.047 ± 0.031 to 0.535 ± 0.148 at 22°C, 0.179 ± 0.143 to 0.458 ± 0.105 at 32°C, and 0.146 ± 0.059 to 0.422 ± 0.069 at 34°C (Fig 4A; Tables S5; all *P* ≤ 0.011). In contrast, females that did not remate showed no significant change (Tables S5; all *P* ≥ 0.198). In *D. melanogaster*, females were allowed to remate after Day 4 (Fig. 1; Fig. 4B). At 34°C, hatch rate increased from 0.538 ± 0.076 to 0.779 ± 0.080 in females that remated (*P* = 0.023), whereas females that did not remate declined from 0.675 ± 0.064 to 0.404 ± 0.066 (Tables S5; *P* = 0.001). These results suggest that access to a new ejaculate can restore or maintain fertility following either heat-induced fertility loss or the natural decline in fertility through time, indicating that the heat-induced fertility loss was likely due to a reduction in functional stored sperm, rather than persistent disruption of oogenesis or female aging.

### Total number of eggs laid per female

Heat exposure also reduced egg production in most species, although responses varied among taxa (Fig. 5). On Day 1, egg production was significantly lower at 32°C in *D. sechellia*, *D. santomea*, *D. yakuba*, and *D. teissieri*, with reductions of approximately 4–12 eggs (32– 67% decrease) relative to 22°C (all *P* ≤ 0.018). In *D. simulans* and *D. melanogaster*, significant reductions occurred only at 34°C (57% and 58%, respectively; Fig. 5; both *P* < 0.001). In contrast, no significant reduction was detected in *D. mauritiana* (*P* = 0.064). Most temperature differences disappeared by Day 4 or Day 8, indicating that the effects of heat on egg production were generally transient. Overall, heat-induced reductions in egg production occurred in fewer species than reductions in egg hatch rate, particularly following exposure to 32°C.

**Fig. 5.**
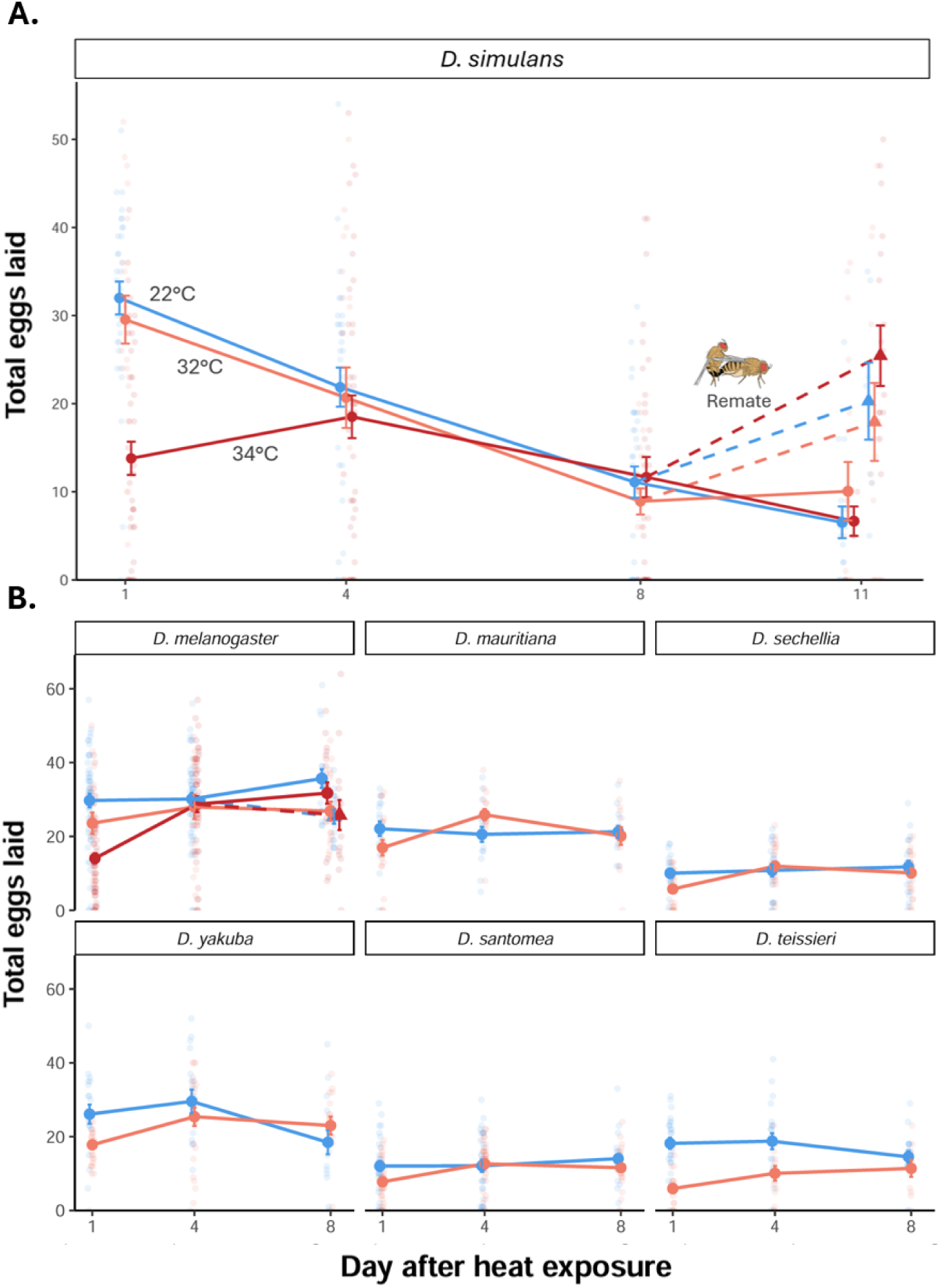
Egg production following heat exposure in *Drosophila*. **(A)** Total number of eggs laid by mated *D. simulans* females exposed to 22°C (blue), 32°C (orange), or 34°C (red). Egg production was assessed on Days 1, 4, and 8 following heat exposure. A subset of females was remated after Day 8 and reassessed on Day 11 (triangles and dashed lines). **(B)** Total number of eggs laid by mated females from six other *Drosophila* species following exposure to 22°C (blue), 32°C (orange), or 34°C (red; *D. melanogaster* only). Points represent mean egg number ± 1 SE. In *D. melanogaster*, triangles and dashed lines indicate females remated after Day 4, whereas circles and solid lines represent females that did not remate.

Remating increased egg production in *D. simulans* (Fig. 5A). Following remating after Day 8, observed mean egg production increased from 6.58 ± 2.15 to 20.29 ± 4.37 eggs at 22°C, from 7.12 ± 2.94 to 17.92 ± 4.42 eggs at 32°C, and from 12.89 ± 3.08 to 25.44 ± 3.44 eggs at 34°C by Day 11 (Fig. 5; all *P* ≤ 0.023). However, remating did not significantly increase egg production in *D. melanogaster* (Fig. 5 B; all *P* ≥ 0.070).

### Stored sperm in the seminal receptacle (SR)

We quantified the number of sperm remaining in the seminal receptacle (SR) in two closely related species that showed distinct patterns of fertility through time (Fig. 6). In *D. simulans*, sperm number declined sharply from Day 1 to Day 4 at all temperatures (Fig. 6A; all *P* ≤ 0.007; Supplementary Table S8) and remained low on Day 8. On Day 4, observed sperm number tended to decrease with increasing temperature, from 33.4 ± 7.2 at 22°C to 22.1 ± 7.6 at 32°C and 16.2 ± 3.3 at 34°C, although temperature differences were not statistically significant (Fig. 6 A ; all pairwise *P* ≥ 0.174). By Day 8, sperm stores were generally low across all temperatures, with many females retaining only single-digit numbers of sperm in the SR (Fig. 6 A).

**Fig. 6.**
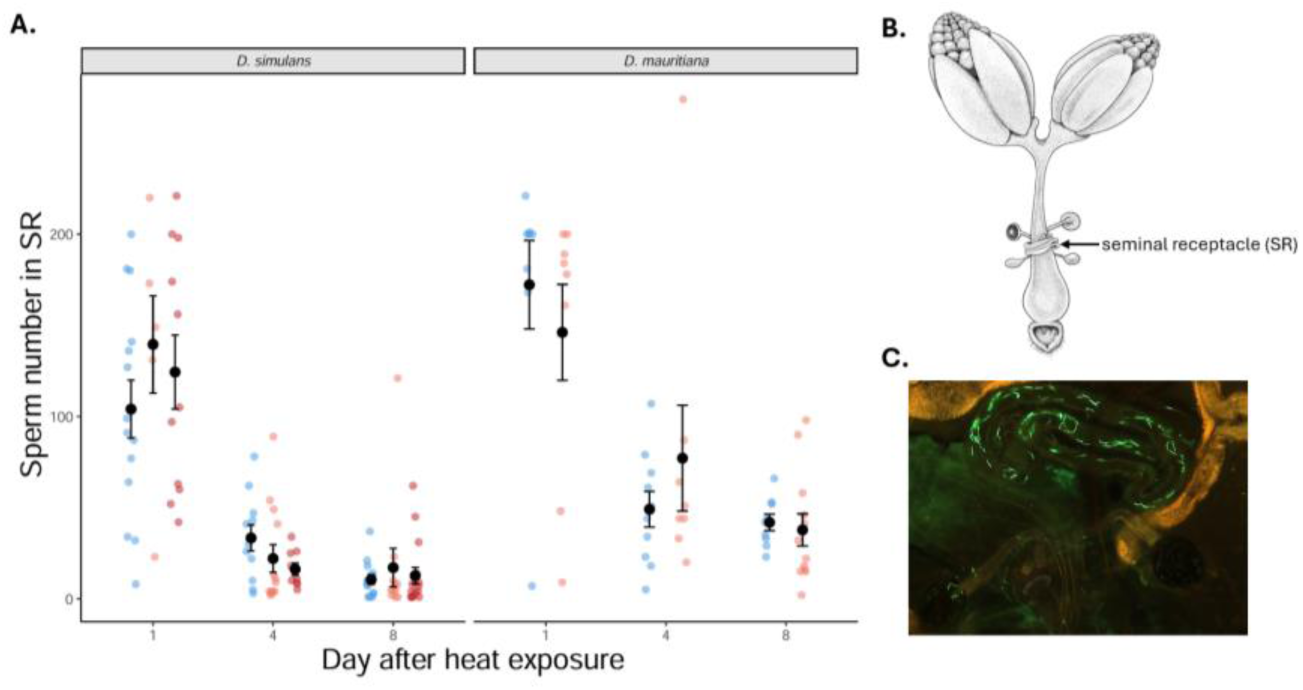
Stored sperm in the seminal receptacle of *D. simulans and D. mauritiana* females. **(A)** Total number of sperm remaining in the seminal receptacle (SR) at different sampling days following exposure to 22°C (blue), 32°C or 34°C (red). Colored points represent individual females, while black points and error bars indicate means ± 1 SE. **(B)** Diagram of the female reproductive tract showing the location of the seminal receptacle (SR) and spermathecae, the two sperm storage organs in female *Drosophila*. **(C)** Fluorescence image of GFP-labeled sperm stored within the seminal receptacle.

*D. mauritiana* also lost sperm through time, with significant declines from Day 1 to Day 8 at both 22 and 32°C (Fig. 6 A; both *P* ≤ 0.025). The two species did not differ significantly in sperm number on Day 1 at either temperature (both *P* ≥ 0.276; Supplementary Table S9), but by Day 8 *D. mauritiana* retained significantly more sperm than *D. simulans* at both 22 and 32°C (both *P* ≤ 0.002). Compared with *D. simulans*, sperm number in *D. mauritiana* remained relatively stable between Days 4 and 8 at both 22 and 32°C (both *P* ≥ 0.178), whereas *D. simulans* showed a further decline over this interval at 22°C (*P* = 0.011). This divergence through time suggests that the species differ in the retention or use of stored sperm rather than in their initial sperm stores or direct response to temperature.

### Association between fertility thermal sensitivity and CTmax

Across species, immediate fertility thermal sensitivity on Day 1 was associated with adult heat tolerance (CTmax; Fig. 7A). Species with higher CTmax values generally showed smaller reductions in hatch rate at 32°C relative to 22°C (adjusted *R*² = 0.566, *P* = 0.031). In contrast, cumulative reproductive loss across Days 1, 4, and 8 showed no significant association with CTmax (Fig. 7B; adjusted *R*² = 0.162, *P* = 0.202). Thus, whole-organism heat tolerance appears to covary with the immediate thermal sensitivity of reproduction, whereas longer-term reproductive consequences might be more strongly shaped by species-specific recovery dynamics.

**Fig. 7.**
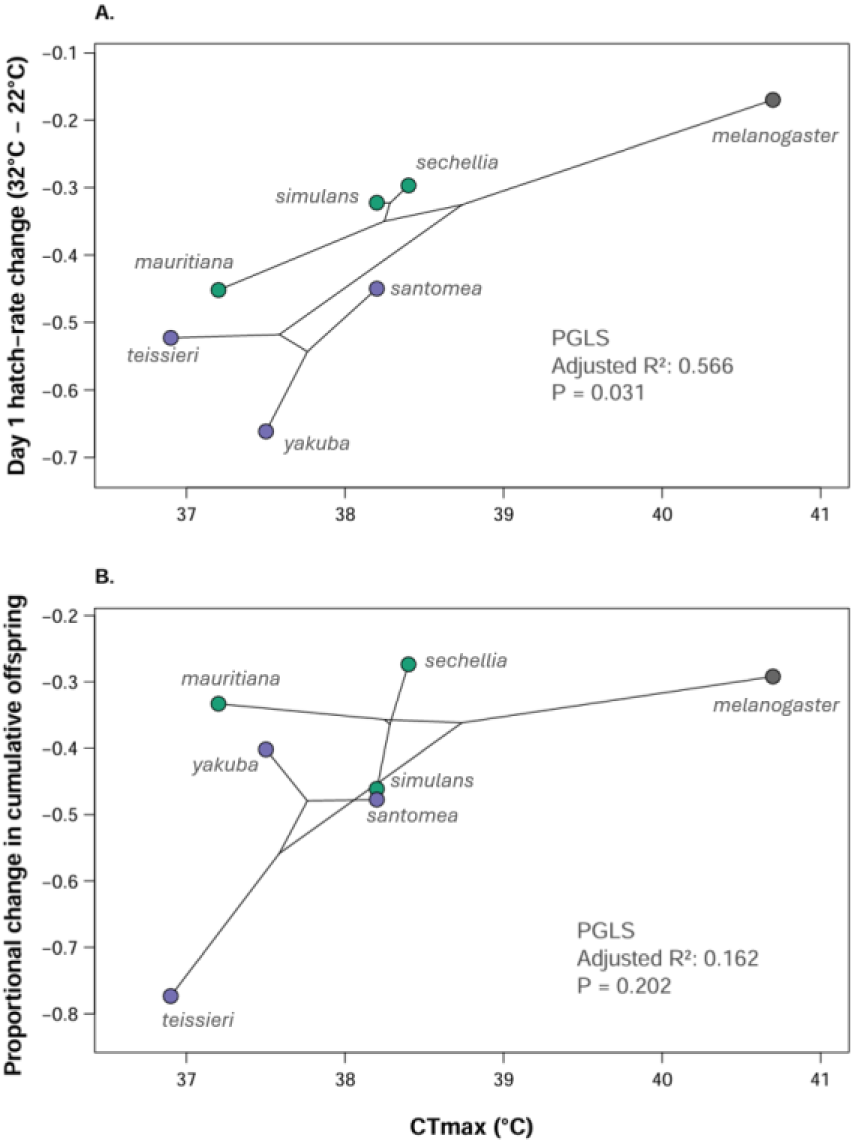
Association between adult heat tolerance and female fertility thermal sensitivity across seven *Drosophila* species. **(A)** Immediate fertility thermal sensitivity, quantified as the difference in Day 1 hatch rate between 32°C and 22°C (32°C – 22°C). More negative values indicate greater immediate fertility loss following heat exposure. **(B)** Cumulative reproductive thermal sensitivity, quantified as the proportional change in cumulative offspring production across Days 1, 4, and 8 between 32°C and 22°C. More negative values indicate greater cumulative reproductive loss. Points represent species, colors indicate phylogenetic groups, and connecting lines show phylogenetic relationships projected into trait space. Associations were evaluated using phylogenetic generalized least squares (PGLS).

## Discussion

Heat-induced fertility loss is widespread among ectotherms, but the initial decline in reproductive performance captures only part of the response to thermal stress. Across seven *Drosophila* species, acute heat exposure consistently reduced fertility, yet the persistence and cumulative consequences of this decline differed substantially among species. Most species recovered within several days, whereas *D. simulans* and *D. melanogaster* showed more persistent effects that were improved by access to a new mating opportunity. Direct sperm counts further showed that *D. simulans* and *D. mauritiana* began with similar sperm abundance but diverged markedly in sperm retention through time. Finally, adult heat tolerance was associated with immediate fertility sensitivity, but not with cumulative reproductive loss. Together, these results suggest that similar immediate fertility losses can be followed by very different recovery trajectories, which may reflect variation in reproductive processes including oocyte turnover, sperm-storage dynamics, and remating. We discuss these findings in turn below.

### Heat-induced fertility loss is widespread across species

Heat exposure reduced egg-hatching success during the first 24 hours following treatment in all seven species, although the magnitude varied substantially (Fig. 4; Tables S5). At 32°C, significant reductions occurred in six species, whereas *D. melanogaster* showed a significant reduction only following exposure to 34°C. Although heat exposure also reduced egg production in most species (Fig. 5), these effects were generally less consistent and weaker than those on hatch rate. For example, in *D. simulans*, egg production declined by only 8% at 32°C relative to 22°C, whereas hatch rate declined by 38%. Similarly, *D. mauritiana* showed no significant reduction in egg production despite an approximately 40% reduction in hatch rate (Fig. 4, 5). Females therefore often continued producing eggs even when relatively few successfully hatched, indicating that heat affected reproductive processes beyond oviposition. These immediate losses in reproductive output were not necessarily offset when fertility subsequently recovered. Although hatch rate recovered within several days in many species (discussed below), cumulative offspring production across the experimental period remained significantly lower following heat exposure in five of the seven species (Fig. 3). A short period of reproductive failure may consequently impose a meaningful fitness cost even when fertility subsequently recovers.

### Some, but not all, species recovered

Despite the widespread Day 1 response, recovery differed strikingly among species. Hatch rate no longer differed between 22 and 32°C by day 4 in five of the seven species (Fig. 4; Supplementary Table S3, S4). Egg production recovered even more consistently, with most species returning to control levels (Fig. 5; Supplementary Table S6, S7). These patterns suggest that much of the initial reproductive impairment in these species was reversible rather than permanent damage to the reproductive system. Several processes could generate this rapid recovery. For example, heat stress can transiently suppress oogenesis and ovulation through endocrine and neural regulation of the female reproductive tract (Meiselman et al., 2018). Oocytes at different stages of development may differ in their sensitivity to heat (Meena et al., 2024; Rivera-Rincón et al., 2024). Because oogenesis is a continuous process in *Drosophila*, heat could disproportionately affect mature or nearly mature oocytes present during exposure, while less developed oocytes that remain relatively unaffected could then mature over the following several days, restoring fertility by Day 4. Species differences in the timing of oocyte development could also influence how quickly fertility recovers following heat exposure (Lavista-Llanos et al., 2014; R’Kha et al., 1997; Rivera-Rincón et al., 2024; Meena et al., 2024; Markow et al., 2009; Horne-Badovinac, 2014). In addition, some *Drosophila* species retain fertilized eggs within the female for longer periods before oviposition. In particular, *D. sechellia* and, *D. yakuba* can retain fertilized eggs and allow substantial embryonic development within the female (Markow et al., 2009). Heat exposure in these species could then affect both developing oocytes and retained early embryos, which may be particularly sensitive to thermal stress (Lockwood et al., 2017; Mikucki et al., 2024; O’Leary et al., 2026).

Recovery without remating also argues against persistent sperm depletion or damage as the primary cause of heat-induced fertility loss in these species, because females regained normal hatching success without receiving new sperm. However, this does not rule out effects on stored sperm; some sperm may have been damaged while sufficient functional sperm remained. Other sperm related processes, including temporary changes in sperm release or utilization, and physiological interactions between sperm, semen and the female reproductive tract, must also be considered.

### Remating restores or maintains fertility in *D. melanogaster* and *D. simulans*

Not all species recovered spontaneously. Females *D. simulans* exposed to elevated temperatures showed little recovery by Day 4 and subsequently experienced an even stronger decline in fertility (Fig. 4 A). In *D. melanogaster* females exposed to 34°C, fertility also failed to recover through Day 8 (Fig. 4B). In both species, however, providing females with a new mating opportunity substantially improved subsequent reproductive performance (Fig. 4). This pattern is consistent with the prediction that, if persistent fertility loss was associated with heat-induced sperm limitation, providing females with a new mating opportunity should restore fertility. Comparable patterns have been reported in other *Drosophilids*. In *Drosophila virilis* and *Zaprionus indianus*, heat exposure after mating caused pronounced fertility loss and remained low until females remated, whereas exposing females to heat before mating had little effect (Walsh et al., 2022).

It is also worth noting that remating affects reproductive function beyond hatching success. In *D. simulans,* females also increased egg production following a new mating opportunity (Fig. 5A), whereas egg production did not significantly increase in *D. melanogaster*. The two species may therefore respond to an additional mating opportunity through somewhat different combinations of reproductive processes. In *D. melanogaster*, improved hatching success without a significant increase in egg production is consistent with the benefit occurring downstream of egg production, potentially through the addition of functional sperm or other ejaculate components affecting fertilization. However, in *D. simulans*, the increase in both egg production and hatch rate suggests a broader reproductive response. For example, mating behavior itself, and seminal-fluid proteins can stimulate ovulation, egg laying, and many other aspects of female reproductive physiology (Pitnick et al., 2009; Avila et al., 2011), so the effect of an additional mating opportunity cannot be attributed to sperm replenishment alone.

### Fertility declines through time and stored sperm may become limiting

Our results also reveal that the ability to maintain fertility differs among species. In *D. simulans*, fertility declined strongly through time even at 22°C, from a mean hatch rate of 0.814 on Day 1 to only 0.100 by Day 8 (Fig. 4A). This decline therefore cannot be attributed solely to heat exposure. Instead, it indicates that reproductive capacity naturally decreases rapidly after initial mating in this species. In contrast, *D. mauritiana* showed a more moderate decline (Fig. 4B; 0.784 ± 0.066 to 0.427 ± 0.093), while *D. santomea* maintained high fertility throughout the same period (0.805 ± 0.052, 0.735 ± 0.063, and 0.899 ± 0.020 on Days 1, 4, and 8, respectively). These differences suggest that species vary in how long fertility can be maintained after mating, potentially through differences in sperm storage and retention that interact with thermal stress to determine how long fertility can be maintained under heat exposure.

Direct sperm counts support this interpretation. *D. simulans* and *D. mauritiana* did not differ significantly in sperm abundance on Day 1, but their trajectories diverged thereafter (Fig. 6A). In *D. simulans*, sperm abundance declined sharply between Days 1 and 4 across all temperatures and remained low by Day 8; at 22°C, sperm abundance declined further between Days 4 and 8, and by Day 8 many females retained only single-digit numbers of sperm (Fig. 6 A). This pattern closely matched the very low hatch rate observed at the same stage of the experiment. In contrast, *D. mauritiana* remained relatively stable sperm abundance between Days 4 and 8, and females retained significantly more sperm than *D. simulans* by Day 8. This pattern points toward species differences in sperm retention or utilization rather than simply differences in the number of sperm initially stored.

Temperature itself, however, had no significant effect on sperm abundance. In *D. simulans*, observed sperm abundance on Day 4 tended to decline with increasing temperature, from 33.4 ± 7.2 sperm at 22°C to 22.1 ± 7.6 at 32°C and 16.2 ± 3.3 at 34°C. However, these differences were far from statistically significant (all *P* ≥ 0.174; Supplementary Table S8). The sperm-count data therefore reveal clear differences in storage dynamics through time, but do not by themselves identify the mechanism responsible for heat-induced fertility loss. However, this result does not rule out heat-induced changes in sperm function, as fluorescent sperm counts provide information on the number of sperm retained in storage but not their viability or fertilizing ability. Females may therefore retain substantial numbers of sperm while experiencing reduced fertility if those sperm have been functionally impaired by heat (McAfee et al., 2020; Sales et al., 2018). Such effects have been demonstrated more directly in other insects. For example, thermal stress can reduce sperm viability within the spermatheca in queen honeybee (McAfee et al., 2020), and compromises stored sperm and subsequent reproductive output in mated red flour beetles (McAfee et al., 2020; Sales et al., 2018).

Nevertheless, when the fertility tracking, remating, and sperm-storage experiments are considered together, sperm availability remains a plausible contributor to the late decline in fertility in *D. simulans*. Stored sperm became increasingly scarce through time as hatch rate simultaneously fell to very low levels, while access to a new ejaculate improved fertility. The fact that fertility also declined rapidly at 22°C suggests that this pattern cannot be explained by direct heat-induced sperm depletion alone, but instead reflects species-specific sperm-storage dynamics acting alongside the acute effects of heat on female. Heat may therefore impose an immediate fertility cost, while the naturally rapid loss or use of stored sperm limits the subsequent ability to maintain reproduction.

### Immediate and cumulative fertility sensitivity show contrasting relationships with adult heat tolerance

The comparative analysis further showed that adult heat tolerance was associated with the immediate, but not longer-term, cumulative reproductive response to heat (Fig. 7). Species with higher CTmax experienced smaller reductions in Day 1 hatch rate following exposure to 32°C (Fig. 7A; *R*² = 0.566, *P* = 0.03), suggesting some covariation between whole-organism and reproductive thermal sensitivity. Such covariation could arise through a shared physiological or genetic basis between these traits, or through thermal coadaptation if both are shaped by similar environmental selection pressures (Angilletta, 2009; Angilletta et al., 2006; Halliday and Blouin-Demers, 2015, 2017). However, this interpretation should remain cautious given the small number of species and the use of a fixed 32°C treatment. Nevertheless, this pattern is broadly consistent with previous comparative work across more *Drosophila* species, where both male and female fertility thermal limits correlate with thermal tolerance and vulnerability to experimental warming (David et al., 2005; van Heerwaarden & Sgrò, 2021; Parratt et al., 2021).

Interestingly, CTmax was not associated with cumulative reproductive loss across Days 1, 4, and 8 (Fig. 7B), suggesting that whole-organism heat tolerance covaries more closely with the immediate thermal sensitivity than with its longer-term consequences. These longer-term outcomes appear to depend more strongly on species-specific recovery dynamics. Indeed, species with similar immediate fertility losses may experience very different cumulative costs depending on what happens after heat exposure. For example, *D. yakuba* showed one of the strongest immediate responses to heat, but hatch rate at 32°C recovered from 0.147 on Day 1 to 0.609 by Day 4. In contrast, *D. simulans* showed a smaller initial decline, yet hatch rate continued to fall from 0.491 on Day 1 to 0.305 on Day 4 and 0.084 by Day 8. Thus, measuring reproductive thermal sensitivity at a single time point may provide only a partial picture of reproductive vulnerability to heat. The contrasting relationships of CTmax with immediate and cumulative fertility sensitivity reveal an interesting new aspect of the link between reproductive and whole-organism thermal physiology and warrant further comparative study.

### Further considerations under climate change: sperm storage, remating, and mating-system evolution

Under current climate-change projections, organisms are expected to experience extreme temperatures more frequently, making it increasingly important to understand how different life stages and components of reproduction respond to thermal stress (Meehl and Tebaldi, 2004; Huey et al., 2012). The association between adult heat tolerance and fertility thermal sensitivity suggests some evolutionary capacity for reproductive heat tolerance, consistent with evidence that warmer-adapted *Drosophila* populations or species often show greater reproductive thermal resilience (van Heerwaarden & Sgrò, 2021; Parratt et al., 2021; Wang and Gunderson, 2022). However, whether reproductive traits can evolve rapidly enough to keep pace with increasingly frequent and intense heat events remains unclear.

At the same time, remating may provide an important behavioral buffer. Our results show that a new mating opportunity can largely rescue fertility in some cases, suggesting that under natural conditions facultative remating could compensate for heat-induced reproductive failure. From an evolutionary perspective, thermal environment, sperm storage dynamics, and mating frequency may therefore interact to shape mating-system evolution. While this remains a relatively new area of study, recent empirical studies support this idea. In *D. pseudoobscura*, females were more likely to remate after mating with heat-stressed, low-fertility males, and remating with fertile males restored normal fecundity (Sutter et al., 2019). Similar results have also been found in *D. subobscura* and flour beetle (*T. castaneum*), suggesting that behavioral polyandry may provide an important form of plastic compensation under increasingly variable thermal environments and may ultimately alter selection on mating systems (Pembury Smith et al., 2025; Vasudeva et al., 2021)

## Conclusion

Consistent with a growing body of evidence, our results reinforce the idea that reproductive failure can occur well before temperatures become lethal and that assessing survival alone may underestimate biological vulnerability to thermal extremes. We also demonstrate that heat-induced fertility loss in mated females is widespread across *Drosophila*, but that the initial decline alone reveals only part of the reproductive response. Similar heat-induced fertility loss can arise through distinct reproductive processes with very different recovery trajectories and consequences through time across species. Together, these findings highlight the value of an integrative understanding of multiple reproductive processes including, but not limited to female reproductive physiology, gametes sensitivity, sperm storage dynamic and mating behavior across taxa. Such an approach will be essential for predicting the resilience of natural populations in a changing world.

## Acknowledgments

We thank all members of the UNL fly group for their assistance, valuable discussions, and feedback throughout this project. We are especially grateful to Dr. Ian Keesey for providing fly stocks and constructive feedback throughout the project and on the manuscript, and to Dr. Javier Torres for his constructive comments on the manuscript. We also thank Rou Rung Chen for the illustrations used in the figures and for helpful discussions. This work was supported by funding from the University of Nebraska–Lincoln to KLM and CDM and by the UNL Program of Excellence in Population Biology Postdoctoral Fellowship to WW-YW.

## Data availability

Data are available in a GitHub repository: https://github.com/wwywanglizardsperm/drosoheat_fertility_data

## Supplementary

**Table S1.**
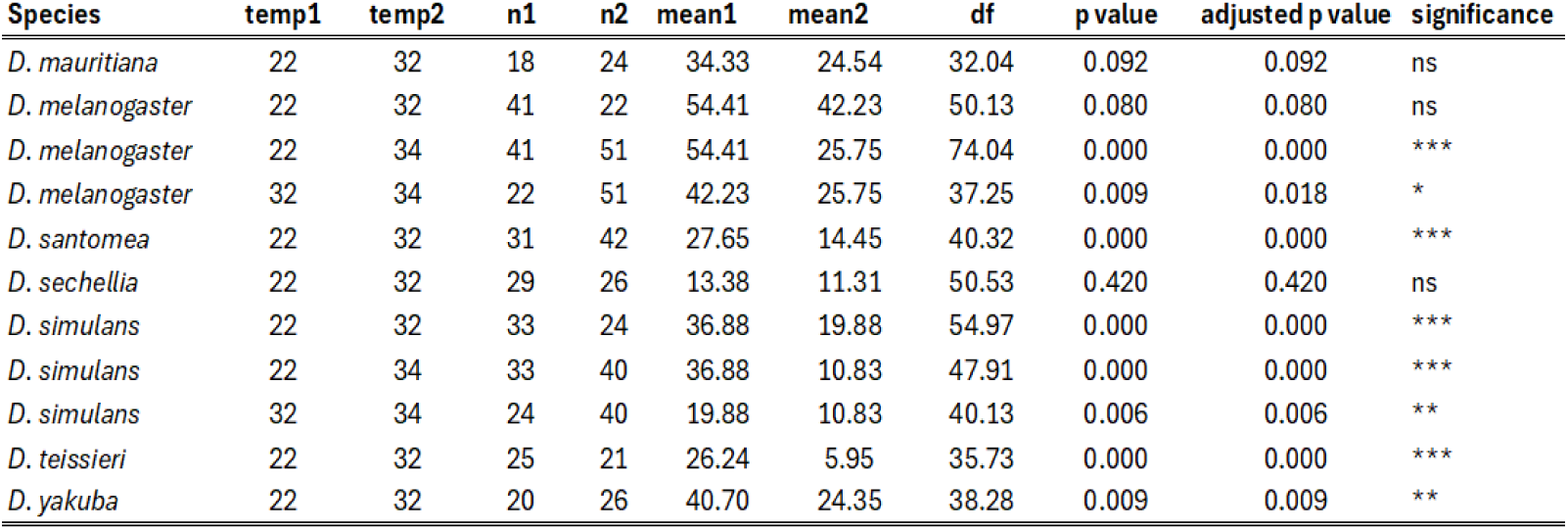
Pairwise comparisons of cumulative reproductive output among temperature treatments across seven *Drosophila* species. Cumulative reproductive output was quantified as the total number of hatched offspring produced per female across Days 1, 4, and 8. Pairwise differences between temperature treatments were evaluated using Welch’s *t*-tests. n1 and n2 indicate sample sizes and mean1 and mean2 indicate mean cumulative reproductive output for the corresponding temperature treatments. *P*-values were adjusted within species using the Holm method to account for multiple comparisons. Significance codes: ns, *P* ≥ 0.05; *, *P* < 0.05; **, *P* < 0.01; ***, *P* < 0.001.

**Table S2.** Hatch rate and egg production across temperature treatments and sampling days in seven *Drosophila* species. Observed hatch rate and total number of eggs laid per female across temperature treatments and sampling days in seven *Drosophila* species. Values are means ± SE, with *n* indicating the number of females contributing to each estimate. Hatch-rate summaries include only observations with at least five scored eggs, whereas all available egg-count observations were retained for egg production. Females were sampled on Days 1, 4, and 8 following thermal exposure. The 34°C treatment was applied only to *D. melanogaster* and *D. simulans*.

| Species | Temp. | Day | n | Mean | SE | n | Mean | SE |
| --- | --- | --- | --- | --- | --- | --- | --- | --- |
|  |  |  | Hatch_rate |  |  | Egg_number |  |  |
| *D. mauritiana* | 22 | 1 | 17 | 0.797 | 0.06 | 18 | 22.11 | 1.95 |
|  | 22 | 4 | 16 | 0.701 | 0.06 | 16 | 20.56 | 2.06 |
|  | 22 | 8 | 14 | 0.427 | 0.09 | 14 | 21.29 | 1.60 |
|  | 32 | 1 | 18 | 0.345 | 0.04 | 22 | 16.95 | 2.10 |
|  | 32 | 4 | 17 | 0.695 | 0.05 | 17 | 25.88 | 1.55 |
|  | 32 | 8 | 13 | 0.447 | 0.09 | 14 | 20.14 | 2.38 |
| *D. melanogaster* | 22 | 1 | 19 | 0.864 | 0.05 | 20 | 30.30 | 3.00 |
|  | 22 | 4 | 18 | 0.805 | 0.05 | 18 | 30.39 | 2.66 |
|  | 22 | 8 | 13 | 0.505 | 0.10 | 13 | 37.38 | 3.24 |
|  | 32 | 1 | 19 | 0.694 | 0.05 | 20 | 23.60 | 2.87 |
|  | 32 | 4 | 18 | 0.791 | 0.05 | 20 | 28.00 | 3.26 |
|  | 32 | 8 | 15 | 0.377 | 0.08 | 15 | 26.87 | 2.59 |
|  | 34 | 1 | 17 | 0.601 | 0.06 | 26 | 12.77 | 2.17 |
|  | 34 | 4 | 16 | 0.585 | 0.10 | 18 | 25.94 | 3.61 |
|  | 34 | 8 | 14 | 0.349 | 0.07 | 14 | 32.93 | 3.70 |
| *D. santomea* | 22 | 1 | 27 | 0.805 | 0.05 | 32 | 12.03 | 1.33 |
|  | 22 | 4 | 19 | 0.735 | 0.06 | 29 | 12.10 | 1.67 |
|  | 22 | 8 | 25 | 0.899 | 0.02 | 26 | 14.04 | 1.13 |
|  | 32 | 1 | 27 | 0.355 | 0.05 | 40 | 7.78 | 0.88 |
|  | 32 | 4 | 30 | 0.675 | 0.04 | 38 | 12.66 | 1.00 |
|  | 32 | 8 | 24 | 0.834 | 0.03 | 27 | 11.59 | 1.05 |
| *D. sechellia* | 22 | 1 | 19 | 0.72 | 0.06 | 22 | 10.05 | 1.06 |
|  | 22 | 4 | 15 | 0.617 | 0.07 | 21 | 10.81 | 1.59 |
|  | 22 | 8 | 17 | 0.276 | 0.08 | 21 | 11.76 | 1.67 |
|  | 32 | 1 | 16 | 0.424 | 0.06 | 26 | 5.77 | 0.92 |
|  | 32 | 4 | 20 | 0.62 | 0.07 | 24 | 11.96 | 1.24 |
|  | 32 | 8 | 19 | 0.303 | 0.07 | 24 | 10.12 | 1.24 |
| *D. simulans* | 22 | 1 | 31 | 0.814 | 0.04 | 32 | 32.00 | 1.86 |
|  | 22 | 4 | 27 | 0.538 | 0.06 | 32 | 21.88 | 2.22 |
|  | 22 | 8 | 19 | 0.1 | 0.04 | 30 | 11.10 | 1.77 |
|  | 32 | 1 | 21 | 0.491 | 0.05 | 22 | 29.55 | 2.72 |
|  | 32 | 4 | 16 | 0.305 | 0.08 | 22 | 20.68 | 3.43 |
|  | 32 | 8 | 15 | 0.084 | 0.04 | 21 | 8.90 | 1.47 |
|  | 34 | 1 | 29 | 0.224 | 0.05 | 39 | 13.79 | 1.89 |
|  | 34 | 4 | 31 | 0.303 | 0.05 | 39 | 18.51 | 2.42 |
|  | 34 | 8 | 19 | 0.161 | 0.05 | 33 | 11.67 | 2.28 |
| *D. teissieri* | 22 | 1 | 24 | 0.749 | 0.04 | 25 | 18.20 | 1.50 |
|  | 22 | 4 | 20 | 0.651 | 0.06 | 22 | 18.77 | 2.18 |
|  | 22 | 8 | 17 | 0.252 | 0.07 | 19 | 14.53 | 1.55 |
|  | 32 | 1 | 11 | 0.226 | 0.05 | 19 | 5.95 | 1.22 |
|  | 32 | 4 | 11 | 0.534 | 0.09 | 14 | 10.07 | 1.98 |
|  | 32 | 8 | 8 | 0.317 | 0.12 | 10 | 11.40 | 2.30 |
| *D. yakuba* | 22 | 1 | 19 | 0.809 | 0.05 | 20 | 26.10 | 2.59 |
|  | 22 | 4 | 17 | 0.583 | 0.07 | 18 | 29.56 | 3.18 |
|  | 22 | 8 | 13 | 0.518 | 0.08 | 15 | 18.47 | 3.24 |
|  | 32 | 1 | 25 | 0.147 | 0.03 | 25 | 17.80 | 0.99 |
|  | 32 | 4 | 21 | 0.609 | 0.06 | 22 | 25.41 | 2.51 |
|  | 32 | 8 | 15 | 0.59 | 0.07 | 17 | 23.00 | 2.42 |

**Table S3.** Pairwise comparisons of hatch rate among temperature treatments within sampling days across seven *Drosophila* species. Pairwise contrasts of hatch rate among temperature treatments on Days 1, 4, and 8 following thermal exposure. Contrasts were obtained from species-specific linear mixed-effects models including temperature, sampling day, and their interaction as fixed effects, with female identity as a random intercept. Estimates represent the difference in estimated marginal mean hatch rate between the first and second temperatures listed in each comparison. *n1* and *n2* indicate the numbers of females contributing to the respective temperature groups. *P*-values were Tukey-adjusted for pairwise comparisons among temperatures within each sampling day. The 34°C treatment was included only for *D. melanogaster* and *D. simulans*. Significance codes: *** *P* < 0.001, ** *P* < 0.01, * *P* < 0.05, ns = not significant.

| Species | Day | Comparison | n1 | n2 | Estimate | SE | df | t | P | Significance |
| --- | --- | --- | --- | --- | --- | --- | --- | --- | --- | --- |
| *D. simulans* | Day 1 | 22 vs 32°C | 31 | 21 | 0.329 | 0.070 | 189.050 | 4.7 | <0.001 | *** |
|  | Day 1 | 22 vs 34 °C | 31 | 29 | 0.578 | 0.064 | 187.620 | 9.041 | <0.001 | *** |
|  | Day 1 | 32 vs 34°C | 21 | 29 | 0.249 | 0.071 | 187.310 | 3.514 | 0.002 | ** |
|  | Day 4 | 22 vs 32°C | 27 | 16 | 0.248 | 0.078 | 196.720 | 3.188 | 0.005 | ** |
|  | Day 4 | 22 vs 34°C | 27 | 31 | 0.235 | 0.066 | 185.890 | 3.578 | 0.001 | ** |
|  | Day 4 | 32 vs 34°C | 16 | 31 | -0.013 | 0.076 | 194.830 | -0.171 | 0.984 | ns |
|  | Day 8 | 22 vs 32°C | 19 | 15 | 0.025 | 0.084 | 198.630 | 0.292 | 0.954 | ns |
|  | Day 8 | 22 vs 34°C | 19 | 19 | -0.055 | 0.079 | 198.520 | -0.696 | 0.766 | ns |
|  | Day 8 | 32 vs 34°C | 15 | 19 | -0.08 | 0.085 | 197.930 | -0.944 | 0.613 | ns |
| *D. sechellia* | Day 1 | 22 vs 32°C | 19 | 16 | 0.298 | 0.098 | 98.580 | 3.032 | 0.003 | ** |
|  | Day 4 | 22 vs 32°C | 15 | 20 | -0.013 | 0.099 | 98.650 | -0.132 | 0.895 | ns |
|  | Day 8 | 22 vs 32°C | 17 | 19 | -0.032 | 0.097 | 98.180 | -0.333 | 0.74 | ns |
| *D. mauritiana* | Day 1 | 22 vs 32°C | 17 | 18 | 0.453 | 0.086 | 74.680 | 5.296 | <0.001 | *** |
|  | Day 4 | 22 vs 32°C | 16 | 17 | 0.002 | 0.088 | 76.350 | 0.028 | 0.978 | ns |
|  | Day 8 | 22 vs 32°C | 14 | 13 | -0.063 | 0.096 | 82.470 | -0.66 | 0.511 | ns |
| *D. santomea* | Day 1 | 22 vs 32°C | 27 | 27 | 0.456 | 0.060 | 139.410 | 7.602 | <0.001 | *** |
|  | Day 4 | 22 vs 32°C | 19 | 30 | 0.056 | 0.064 | 143.460 | 0.873 | 0.384 | ns |
|  | Day 8 | 22 vs 32°C | 25 | 24 | 0.012 | 0.063 | 142.300 | 0.192 | 0.848 | ns |
| *D. melanogaster* | Day 1 | 22 vs 32°C | 19 | 19 | 0.171 | 0.089 | 117.570 | 1.917 | 0.138 | ns |
|  | Day 1 | 22 vs 34°C | 19 | 17 | 0.265 | 0.091 | 124.550 | 2.915 | 0.012 | * |
|  | Day 1 | 32 vs 34°C | 19 | 17 | 0.094 | 0.090 | 129.010 | 1.04 | 0.553 | ns |
|  | Day 4 | 22 vs 32°C | 18 | 18 | 0.023 | 0.091 | 121.940 | 0.248 | 0.967 | ns |
|  | Day 4 | 22 vs 34°C | 18 | 16 | 0.217 | 0.093 | 125.600 | 2.321 | 0.057 | ns |
|  | Day 4 | 32 vs 34°C | 18 | 16 | 0.194 | 0.093 | 131.740 | 2.095 | 0.095 | ns |
|  | Day 8 | 22 vs 32°C | 13 | 15 | 0.138 | 0.102 | 132.350 | 1.356 | 0.367 | ns |
|  | Day 8 | 22 vs 34°C | 13 | 14 | 0.175 | 0.103 | 135.290 | 1.695 | 0.211 | ns |
|  | Day 8 | 32 vs 34°C | 15 | 14 | 0.036 | 0.099 | 135.930 | 0.367 | 0.928 | ns |
| *D. yakuba* | Day 1 | 22 vs 32°C | 19 | 25 | 0.658 | 0.073 | 93.090 | 9.064 | <0.001 | *** |
|  | Day 4 | 22 vs 32°C | 17 | 21 | -0.032 | 0.078 | 96.490 | -0.407 | 0.685 | ns |
|  | Day 8 | 22 vs 32°C | 13 | 15 | -0.074 | 0.089 | 102.440 | -0.829 | 0.409 | ns |
| *D. teissieri* | Day 1 | 22 vs 32°C | 24 | 11 | 0.519 | 0.093 | 84.810 | 5.586 | <0.001 | *** |
|  | Day 4 | 22 vs 32°C | 20 | 11 | 0.113 | 0.096 | 84.800 | 1.177 | 0.242 | ns |
|  | Day 8 | 22 vs 32°C | 17 | 8 | -0.063 | 0.110 | 84.970 | -0.574 | 0.568 | ns |

**Table S4.**
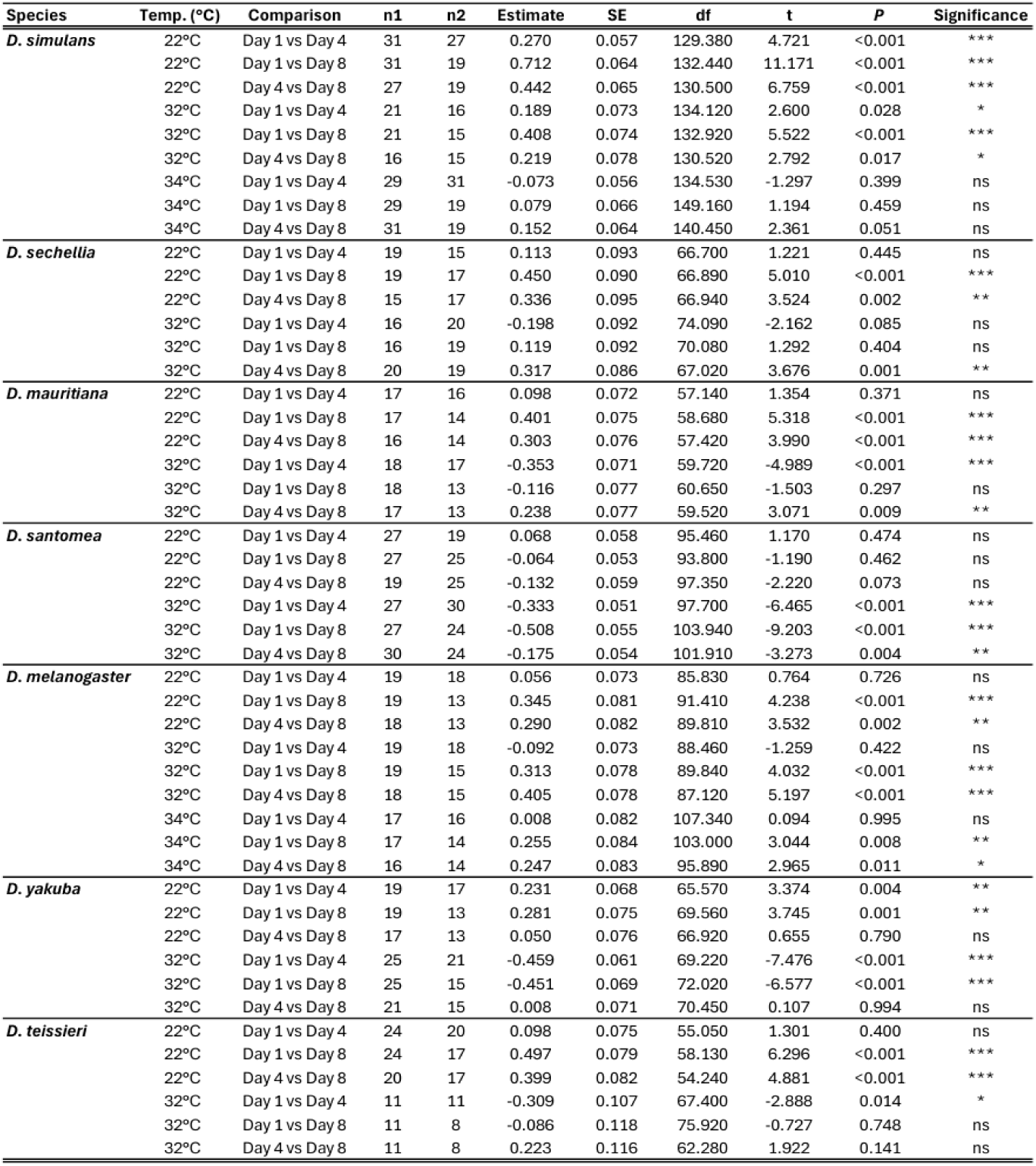
Pairwise comparisons of hatch rate among sampling days within temperature treatments across seven *Drosophila* species. Pairwise contrasts of hatch rate among Days 1, 4, and 8 within each temperature treatment. Contrasts were obtained from species-specific linear mixed-effects models including temperature, sampling day, and their interaction as fixed effects, with female identity as a random intercept. Estimates represent the difference in estimated marginal mean hatch rate between the first and second days listed in each comparison; positive estimates indicate higher hatch rate on the first day and negative estimates indicate higher hatch rate on the second day. *n1* and *n2* indicate the numbers of females contributing to the respective sampling days. *P*-values were Tukey-adjusted for pairwise comparisons among days within each temperature treatment. Hatch-rate analyses included only observations with at least five scored eggs. The 34°C treatment was included only for *D. melanogaster* and *D. simulans*. Significance codes: *** *P* < 0.001, ** *P* < 0.01, * *P* < 0.05, ns = not significant.

**Table S5.** Changes in hatch rate and egg production following remating in *D. melanogaster* and *D. simulans*. Changes in hatch rate and egg production in remated and non-remated females of *D. melanogaster* and *D. simulans*. Females of *D. melanogaster* were given a remating opportunity after Day 4 and responses were compared between Days 4 and 8, whereas *D. simulans* females were given a remating opportunity after Day 8 and responses were compared between Days 8 and 11. Mean 1 and Mean 2 are observed means ± SE for the first and second sampling days listed in each comparison, respectively, and *n1* and *n2* indicate the corresponding sample sizes. Statistical contrasts were obtained from species-specific linear mixed-effects models including temperature, sampling day, remating status, and their interactions, with female identity as a random intercept. Hatch-rate analyses included only observations with at least five scored eggs. Significance codes: *** *P* < 0.001, ** *P* < 0.01, * *P* < 0.05, ns = not significant.

| Hatch rate |  |  |  |  |  |  |  |  |  |  |  |  |  |
| --- | --- | --- | --- | --- | --- | --- | --- | --- | --- | --- | --- | --- | --- |
| Species | Temp. | Mating_status | Comparison | n1 | Mean_1 | SE_1 | n2 | Mean_2 | SE_2 | df | t | P | Significance |
| *D. melanogaster* | 22°C | Not remated | Day 4 vs 8 | 30 | 0.781 | 0.046 | 18 | 0.541 | 0.078 | 83.25 | 2.933 | 0.004 | ** |
|  | 34°C | Not remated | Day 4 vs 8 | 27 | 0.675 | 0.064 | 20 | 0.404 | 0.066 | 82.55 | 3.31 | 0.001 | ** |
|  | 22°C | Remated | Day 4 vs 8 | 14 | 0.627 | 0.081 | 14 | 0.791 | 0.05 | 64.1 | -1.578 | 0.12 | ns |
|  | 34°C | Remated | Day 4 vs 8 | 14 | 0.538 | 0.076 | 14 | 0.779 | 0.08 | 64.1 | -2.329 | 0.023 | * |
| *D. simulans* | 22°C | Not remated | Day 8 vs 11 | 13 | 0.124 | 0.049 | 9 | 0.081 | 0.074 | 56.05 | 0.741 | 0.462 | ns |
|  | 32°C | Not remated | Day 8 vs 11 | 11 | 0.05 | 0.025 | 8 | 0.186 | 0.112 | 59.93 | -1.301 | 0.198 | ns |
|  | 34°C | Not remated | Day 8 vs 11 | 6 | 0.193 | 0.065 | 3 | 0.067 | 0.067 | 88.52 | 0.74 | 0.461 | ns |
|  | 22°C | Remated | Day 8 vs 11 | 6 | 0.047 | 0.031 | 7 | 0.535 | 0.148 | 43.33 | -4.452 | <0.001 | *** |
|  | 32°C | Remated | Day 8 vs 11 | 4 | 0.179 | 0.143 | 10 | 0.458 | 0.105 | 52.42 | -2.647 | 0.011 | * |
|  | 34°C | Remated | Day 8 vs 11 | 13 | 0.146 | 0.059 | 16 | 0.422 | 0.069 | 31.81 | -3.51 | 0.001 | ** |
| Egg number |  |  |  |  |  |  |  |  |  |  |  |  |  |
| Species | Temp. | Mating_status | Comparison | n1 | Mean_1 | SE_1 | n2 | Mean_2 | SE_2 | df | t | P | Significance |
| *D. melanogaster* | 22°C | Not remated | Day 4 vs 8 | 32 | 29.22 | 2.17 | 18 | 35.67 | 2.54 | 79.55 | -2.028 | 0.046 | * |
|  | 34°C | Not remated | Day 4 vs 8 | 31 | 27.23 | 2.83 | 20 | 31.75 | 2.90 | 82.91 | -0.717 | 0.476 | ns |
|  | 22°C | Remated | Day 4 vs 8 | 14 | 32.29 | 2.49 | 14 | 25.36 | 1.90 | 63.14 | 1.844 | 0.070 | ns |
|  | 34°C | Remated | Day 4 vs 8 | 15 | 31.87 | 2.91 | 15 | 25.8 | 4.07 | 63.14 | 1.672 | 0.100 | ns |
| *D. simulans* | 22°C | Not remated | Day 8 vs 11 | 18 | 14.11 | 2.37 | 17 | 6.53 | 1.80 | 94.82 | 2.063 | 0.042 | * |
|  | 32°C | Not remated | Day 8 vs 11 | 13 | 10 | 1.58 | 17 | 10.06 | 3.32 | 90.29 | -0.004 | 0.997 | ns |
|  | 34°C | Not remated | Day 8 vs 11 | 15 | 10.2 | 3.46 | 3 | 6.67 | 1.67 | 144.7 | 0.44 | 0.660 | ns |
|  | 22°C | Remated | Day 8 vs 11 | 12 | 6.58 | 2.15 | 7 | 20.29 | 4.37 | 76.19 | -2.518 | 0.014 | * |
|  | 32°C | Remated | Day 8 vs 11 | 8 | 7.12 | 2.94 | 13 | 17.92 | 4.42 | 83.78 | -2.316 | 0.023 | * |
|  | 34°C | Remated | Day 8 vs 11 | 18 | 12.89 | 3.08 | 18 | 25.44 | 3.44 | 59.9 | -3.383 | 0.001 | ** |

**Table S6.**
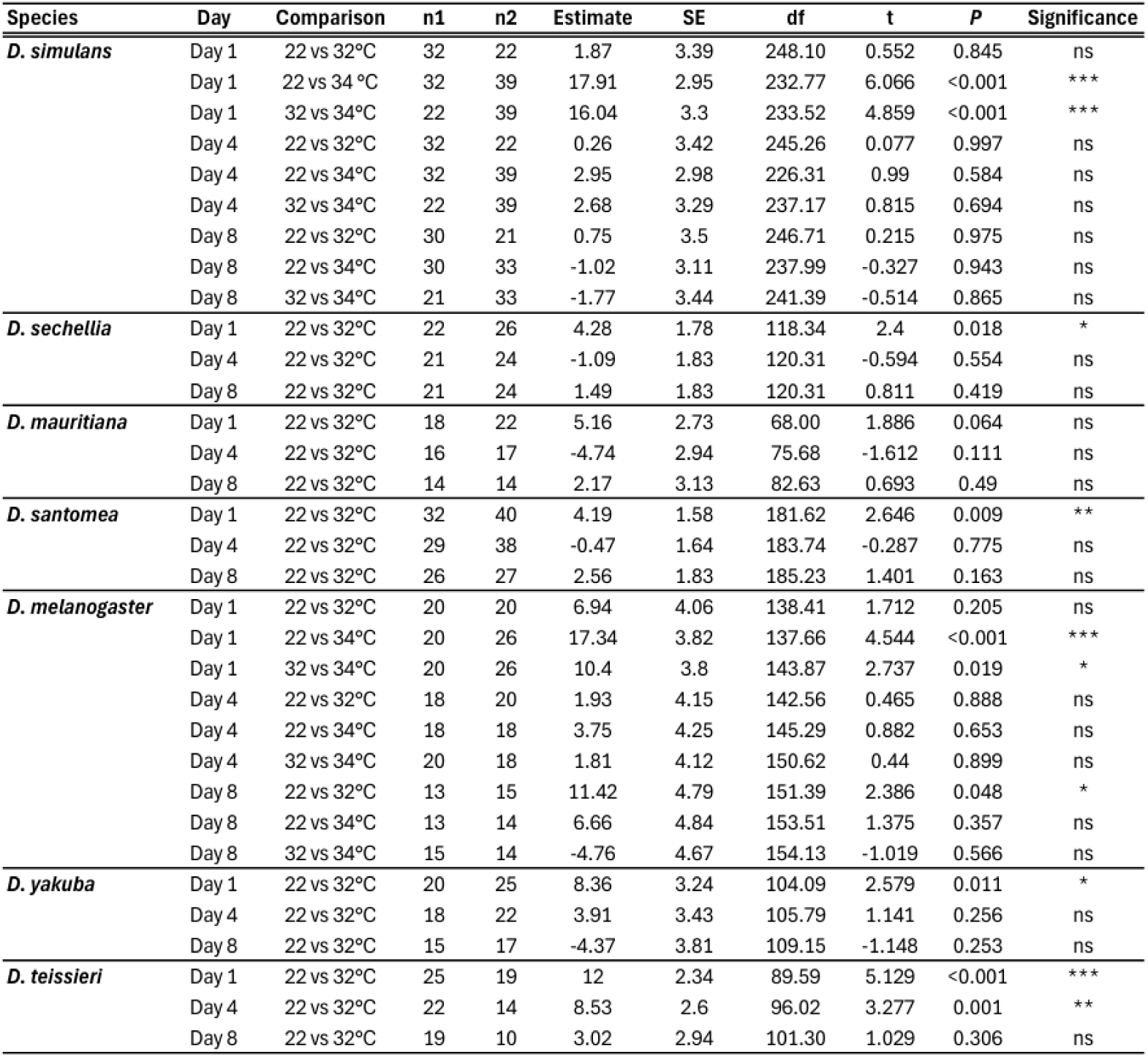
Pairwise comparisons of egg production among temperature treatments within sampling days across seven *Drosophila* species. Pairwise contrasts of total number of eggs laid per female among temperature treatments on Days 1, 4, and 8 following thermal exposure. Contrasts were obtained from species-specific linear mixed-effects models including temperature, sampling day, and their interaction as fixed effects, with female identity as a random intercept. Estimates represent the difference in estimated marginal mean egg production between the first and second temperatures listed in each comparison; positive estimates indicate higher egg production at the first temperature and negative estimates indicate higher egg production at the second temperature. *n1* and *n2* indicate the numbers of females contributing to the respective temperature groups. *P*-values were Tukey-adjusted for pairwise comparisons among temperatures within each sampling day. The 34°C treatment was included only for *D. melanogaster* and *D. simulans*. Significance codes: *** *P* < 0.001, ** *P* < 0.01, * *P* < 0.05, ns = not significant.

**Table S7.**
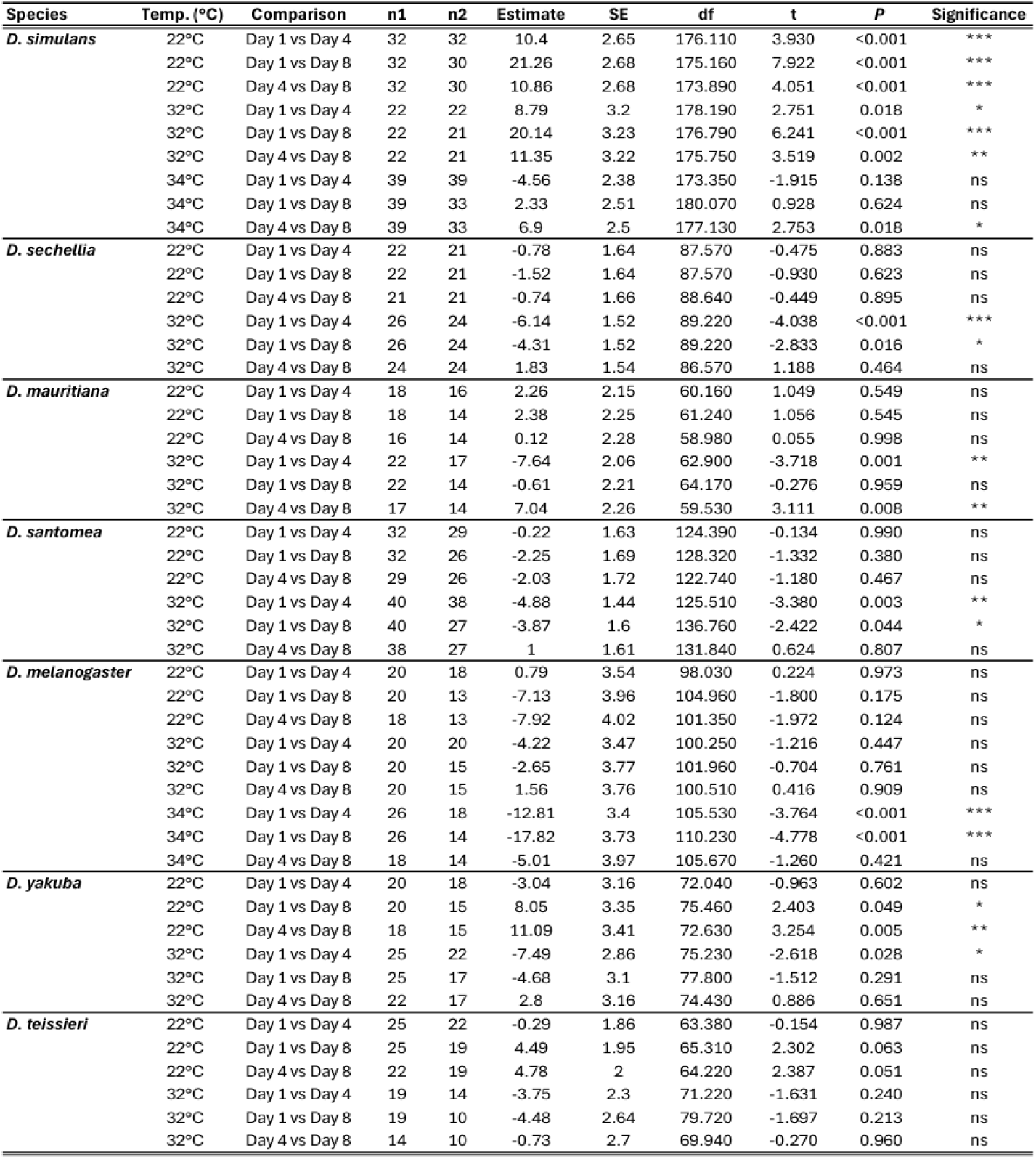
Pairwise comparisons of egg production among sampling days within temperature treatments across seven *Drosophila* species. Pairwise contrasts of total number of eggs laid per female among Days 1, 4, and 8 within each temperature treatment. Contrasts were obtained from species-specific linear mixed-effects models including temperature, sampling day, and their interaction as fixed effects, with female identity as a random intercept. Estimates represent the difference in estimated marginal mean egg production between the first and second days listed in each comparison; positive estimates indicate higher egg production on the first day and negative estimates indicate higher egg production on the second day. *n1* and *n2* indicate the numbers of females contributing to the respective sampling days. *P*-values were Tukey-adjusted for pairwise comparisons among days within each temperature treatment. The 34°C treatment was included only for *D. melanogaster* and *D. simulans*. Significance codes: *** *P* < 0.001, ** *P* < 0.01, * *P* < 0.05, ns = not significant.

**Table S8.**
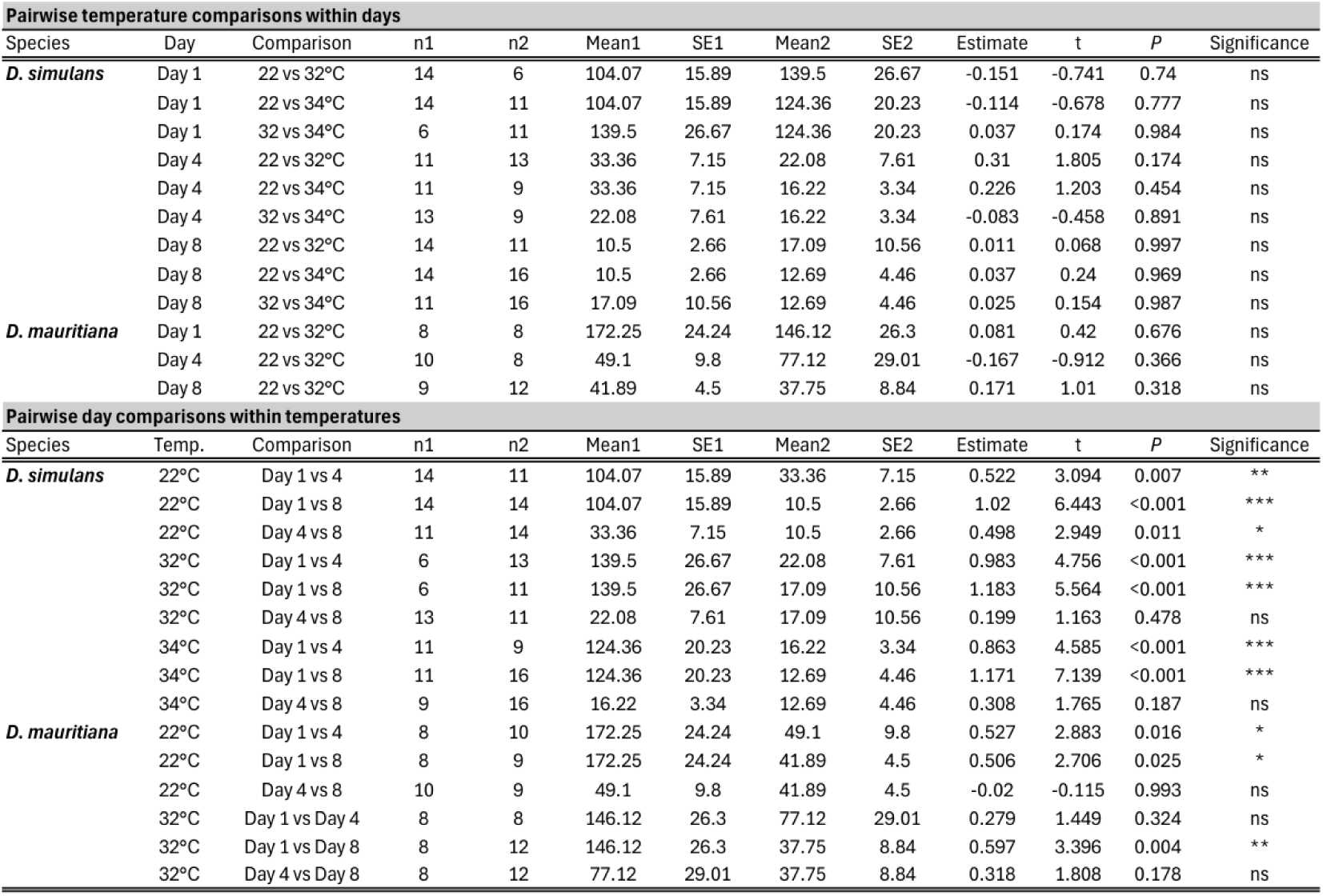
Pairwise comparisons of seminal receptacle sperm abundance across temperature treatments and sampling days in *Drosophila simulans* and *D. mauritiana*. The upper section shows pairwise temperature comparisons within each sampling day, and the lower section shows pairwise day comparisons within each temperature treatment. Mean1 and Mean2 are observed sperm counts presented as mean ± SE, whereas estimates, *t*-statistics, and *P*-values are based on linear models of log10- transformed sperm abundance. Pairwise comparisons were obtained using estimated marginal means with Tukey-adjusted *P*-values. Significance codes: *** *P* < 0.001, ** *P* < 0.01, * *P* < 0.05, ns = not significant.

**Table S9.** Pairwise comparisons of seminal receptacle sperm abundance between *Drosophila simulans* and *D. mauritiana*. Mean_sim. and Mean_mau. are mean of observed sperm counts presented with their corresponding SEs and sample sizes (*n*). Estimates, *t*-statistics, and *P*-values are based on a linear model of log10-transformed sperm abundance including species, temperature, sampling day, and interactions. Estimates represent *D. simulans* − *D. mauritiana*. Significance codes: *** *P* < 0.001, ** *P* < 0.01, * *P* < 0.05, ns = not significant.

| Temp. | Day | Comparison | n_sim. | Mean_sim. | SE_sim. | n_mau. | Mean_mau. | SE_mau. | Estimate | t | P | Significance |
| --- | --- | --- | --- | --- | --- | --- | --- | --- | --- | --- | --- | --- |
| 22°C | Day 1 | D. sim. vs D. mau. | 14 | 104.070 | 15.89 | 8 | 172.250 | 24.24 | -0.201 | -1.094 | 0.276 | ns |
| 32°C | Day 1 | D. sim. vs D. mau. | 6 | 139.500 | 26.67 | 8 | 146.120 | 26.30 | 0.031 | 0.138 | 0.891 | ns |
| 22°C | Day 4 | D. sim. vs D. mau. | 11 | 33.360 | 7.15 | 10 | 49.100 | 9.80 | -0.197 | -1.085 | 0.28 | ns |
| 32°C | Day 4 | D. sim. vs D. mau. | 13 | 22.080 | 7.61 | 8 | 77.120 | 29.01 | -0.673 | -3.607 | <0.001 | *** |
| 22°C | Day 8 | D. sim. vs D. mau. | 14 | 10.500 | 2.66 | 9 | 41.890 | 4.50 | -0.715 | -4.029 | <0.001 | *** |
| 32°C | Day 8 | D. sim. vs D. mau. | 11 | 17.090 | 10.56 | 12 | 37.750 | 8.84 | -0.555 | -3.201 | 0.002 | ** |

## References

Angilletta, M. J., Jr. (2009). Thermal adaptation: A theoretical and empirical synthesis. Oxford University Press. 10.1093/acprof:oso/9780198570875.001.1

Angilletta, M. J., Jr., Bennett, A. F., Guderley, H., Navas, C. A., Seebacher, F., & Wilson, R. S. (2006). Coadaptation: a unifying principle in evolutionary thermal biology. Physiological and biochemical zoology, 79(2), 282–294. 10.1086/499990

Avila, F. W., Sirot, L. K., LaFlamme, B. A., Rubinstein, C. D., & Wolfner, M. F. (2011). Insect seminal fluid proteins: Identification and function. Annual Review of Entomology, 56, 21–40. 10.1146/annurev-ento-120709-144823

Bates, D., Mächler, M., Bolker, B., & Walker, S. (2015). Fitting linear mixed-effects models using lme4. Journal of Statistical Software, 67(1), 1–48. 10.18637/jss.v067.i01

Bundgaard, J., & Barker, J. S. F. (2017). Genetic variation for resistance to high temperature stress of mature sperm – a study in *Drosophila*. PLOS ONE, 12(3), e0173990. 10.1371/journal.pone.0173990

David, J. R., Araripe, L. O., Chakir, M., Legout, H., Lemos, B., Pétavy, G., Rohmer, C., Joly, D., & Moreteau, B. (2005). Male sterility at extreme temperatures: A significant but neglected phenomenon for understanding *Drosophila* climatic adaptations. Journal of Evolutionary Biology, 18, 838–846. 10.1111/j.1420-9101.2005.00914.x

Dougherty, L. R., Frost, F., Maenpaa, M. I., Rowe, M., Cole, B. J., Vasudeva, R., Pottier, P., Schultner, E., Macartney, E. L., Lindenbaum, I., Smith, J. L., Carazo, P., Graziano, M., Weaving, H., Canal Domenech, B., Berger, D., Meena, A., Bishop, T. R., Noble, D. W. A., … Price, T. A. R. (2024). A systematic map of studies testing the relationship between temperature and animal reproduction. Ecological Solutions and Evidence, 5(1), e12303. 10.1002/2688-8319.12303

Halliday, W. D., & Blouin-Demers, G. (2015). A stringent test of the thermal coadaptation hypothesis in flour beetles. J. Therm. Biol., 52, 108*–*116. 10.1016/j.jtherbio.2015.06.002

Halliday, W. D., & Blouin-Demers, G. (2017). A test of the thermal coadaptation hypothesis with ultimate measures of fitness in flour beetles. J. Therm. Biol., 69, 206–212. 10.1016/j.jtherbio.2017.07.017

Horne-Badovinac, S. (2014). The *Drosophila* egg chamber—A new spin on how tissues elongate. Integrative and Comparative Biology, 54(4), 667–676. 10.1093/icb/icu067

Huey, R. B., Kearney, M. R., Krockenberger, A., Holtum, J. A. M., Jess, M., & Williams, S. E. (2012). Predicting organismal vulnerability to climate warming: Roles of behaviour, physiology and adaptation. Philosophical Transactions of the Royal Society B: Biological Sciences, 367(1596), 1665–1679. 10.1098/rstb.2012.0005

Iossa, G. (2019). Sex-specific differences in thermal fertility limits. Trends in Ecology & Evolution, 34(6), 490–492. 10.1016/j.tree.2019.02.016

Kuznetsova, A., Brockhoff, P. B., & Christensen, R. H. B. (2017). lmerTest package: Tests in linear mixed effects models. Journal of Statistical Software, 82(13), 1–26. 10.18637/jss.v082.i13

Lavista-Llanos, S., Svatoš, A., Kai, M., Riemensperger, T., Birman, S., Stensmyr, M. C., & Hansson, B. S. (2014). Dopamine drives *Drosophila sechellia* adaptation to its toxic host. eLife, 3, e03785. 10.7554/eLife.03785

Lenth, R. V. (2024). emmeans: Estimated marginal means, aka least-squares means. R package. https://CRAN.R-project.org/package=emmeans

Lockwood, B. L., Gupta, T., & Scavotto, R. (2018). Disparate patterns of thermal adaptation between life stages in temperate vs. tropical *Drosophila melanogaster*. Journal of Evolutionary Biology, 31(2), 323–331. 10.1111/jeb.13234

Lockwood, B. L., Julick, C. R., & Montooth, K. L. (2017). Maternal loading of a small heat shock protein increases embryo thermal tolerance in *Drosophila melanogaster*. Journal of Experimental Biology, 220(23), 4492–4501. 10.1242/jeb.164848

Markow, T. A., Beall, S., & Matzkin, L. M. (2009). Egg size, embryonic development time and ovoviviparity in *Drosophila* species. Journal of Evolutionary Biology, 22(2), 430–434. 10.1111/j.1420-9101.2008.01649.x

McAfee, A., Chapman, A., Higo, H., Underwood, R., Milone, J., Foster, L. J., Guarna, M. M., Tarpy, D. R., & Pettis, J. S. (2020). Vulnerability of honey bee queens to heat-induced loss of fertility. Nature Sustainability, 3, 367–376. 10.1038/s41893-020-0493-x

Meehl, G. A. and Tebaldi, C. (2004). More intense, more frequent, and longer lasting heat waves in the 21st century. Science, 305, 994–997.

Meena, A., De Nardo, A. N., Maggu, K., Sbilordo, S. H., Roy, J., Snook, R. R., & Lüpold, S. (2024). Fertility loss and recovery dynamics after repeated heat stress across life stages in male *Drosophila melanogaster*: patterns and processes. Royal Society open science, 11(10), 241082. 10.1098/rsos.241082

Meena, A., Maggu, K., De Nardo, A. N., Kovalov, V., Eggs, B., & Lüpold, S. (2025). Stage-specific and cumulative effects of heat stress on male and female reproductive performance in *Drosophila melanogaster*. Journal of Thermal Biology, 132, 104213. 10.1016/j.jtherbio.2025.104213

Meena, A., Maggu, K., De Nardo, A. N., Sbilordo, S. H., Eggs, B., Al Toma Sho, R., & Lüpold, S. (2024). Life stage-specific effects of heat stress on spermatogenesis and oogenesis in *Drosophila melanogaster*. Journal of Thermal Biology, 125, 104001. 10.1016/j.jtherbio.2024.104001

Meiselman, M. R., Kingan, T. G., & Adams, M. E. (2018). Stress-induced reproductive arrest in *Drosophila* occurs through ETH deficiency-mediated suppression of oogenesis and ovulation. BMC Biology, 16, 18. 10.1186/s12915-018-0484-9

Mikucki, E. E., O’Leary, T. S., & Lockwood, B. L. (2024). Heat tolerance, oxidative stress response tuning and robust gene activation in early-stage *Drosophila melanogaster* embryos. Proceedings of the Royal Society B: Biological Sciences, 291(2029), 20240973. 10.1098/rspb.2024.0973

O’Leary, T. S., Mikucki, E. E., Tangwancharoen, S., Boyd, J. R., Helms Cahan, S., Frietze, S., & Lockwood, B. L. (2026). Single-nucleus multiome sequencing reveals the molecular basis of thermal acclimation in Drosophila melanogaster embryos. Development (Cambridge, England), 153(18), dev205729. 10.1242/dev.205729

Parratt, S. R., Walsh, B. S., Metelmann, S., White, N., Manser, A., Bretman, A. J., Hoffmann, A. A., Snook, R. R., & Price, T. A. R. (2021). Temperatures that sterilize males better match global species distributions than lethal temperatures. Nature Climate Change, 11, 481–484. 10.1038/s41558-021-01047-0

Pembury Smith, M. Q. R., Latkova, L., & Snook, R. R. (2025). Facultative polyandry under heat stress and the evolutionary potential for climate-driven shifts in mating systems. Heredity, 134, 596–604. 10.1038/s41437-025-00795-5

Pitnick, S., Wolfner, M. F., & Suarez, S. S. (2009). Ejaculate-female and sperm-female interactions. In T. R. Birkhead, D. J. Hosken, & S. Pitnick (Eds.), Sperm Biology: An Evolutionary Perspective (pp. 247–304). Academic Press. 10.1016/B978-0-12-372568-4.00007-0

R Core Team. (2025). R: A language and environment for statistical computing. R Foundation for Statistical Computing, Vienna, Austria. https://www.R-project.org/

R’Kha, S., Moreteau, B., Coyne, J. A., & David, J. R. (1997). Evolution of a lesser fitness trait: Egg production in the specialist *Drosophila sechellia*. Genetical Research, 69(1), 17–23. 10.1017/S0016672396002546

Rivera-Rincón, N., Altindag, U. H., Amin, R., Graze, R. M., Appel, A. G., & Stevison, L. S. (2024). A comparison of thermal stress response between *Drosophila melanogaster* and *Drosophila pseudoobscura* reveals differences between species and sexes. Journal of Insect Physiology, 153, 104616. 10.1016/j.jinsphys.2024.104616

Sales, K., Vasudeva, R., Dickinson, M. E., Godwin, J. L., Lumley, A. J., Michalczyk, Ł., Hebberecht, L., Thomas, P., Franco, A., & Gage, M. J. G. (2018). Experimental heatwaves compromise sperm function and cause transgenerational damage in a model insect. Nature Communications, 9, 4771. 10.1038/s41467-018-07273-z

Snook, R. R., Bretman, A., Dougherty, L. R., & Fricke, C. (2026). The consequences of rising temperatures for animal fertility. Nature Reviews Biodiversity, 2, 225–241. 10.1038/s44358-026-00142-4

Sutter, A., Travers, L. M., Oku, K., Delaney, K. L., Store, S. J., Price, T. A. R., & Wedell, N. (2019). Flexible polyandry in female flies is an adaptive response to infertile males. Behavioral Ecology, 30(6), 1715–1724. 10.1093/beheco/arz140

van Heerwaarden, B., & Sgrò, C. M. (2021). Male fertility thermal limits predict vulnerability to climate warming. Nature Communications, 12, 2214. 10.1038/s41467-021-22546-w

Vasudeva, R., Dickinson, M., Sutter, A., Powell, S., Sales, K., & Gage, M. J. G. (2021). Facultative polyandry protects females from compromised male fertility caused by heatwave conditions. Animal Behaviour, 178, 37–48. 10.1016/j.anbehav.2021.05.016

Walsh, B. S., Parratt, S. R., Hoffmann, A. A., Atkinson, D., Snook, R. R., Bretman, A., & Price, T. A. R. (2019). The impact of climate change on fertility. Trends in Ecology & Evolution, 34(3), 249–259. 10.1016/j.tree.2018.12.002

Walsh, B. S., Parratt, S. R., Snook, R. R., Bretman, A., Atkinson, D., & Price, T. A. R. (2022). Female fruit flies cannot protect stored sperm from high temperature damage. Journal of Thermal Biology, 105, 103209. 10.1016/j.jtherbio.2022.103209

Wang, W. W.-Y., & Gunderson, A. R. (2022). The physiological and evolutionary ecology of sperm thermal performance. Frontiers in Physiology, 13, 754830. 10.3389/fphys.2022.754830

